# The transfer of antibiotic resistance genes between evolutionary distant bacteria

**DOI:** 10.1101/2024.10.22.619579

**Authors:** Marcos Parras-Moltó, David Lund, Stefan Ebmeyer, DG Joakim Larsson, Anna Johnning, Erik Kristiansson

## Abstract

Infections from antibiotic-resistant bacteria threaten human health globally. Resistance is often caused by mobile antibiotic resistance genes (ARGs) shared horizontally between bacterial genomes. Many ARGs originate from environmental and commensal bacteria and are transferred between divergent bacterial hosts before they reach pathogens. This process remains, however, poorly understood, which complicates the development of countermeasures that reduce the spread of ARGs. In this study, we aimed to systematically analyze the ARGs transferred between the most evolutionary distant bacteria, here defined based on their phylum. We implemented an algorithm that identified inter-phyla transfers (IPTs) by combining ARG-specific phylogenetic trees with the taxonomy of the bacterial hosts. From the analysis of almost 1 million resistance genes identified in >400,000 bacterial genomes, we identified 661 IPTs, which included transfers between all major bacterial phyla. The frequency of IPTs varies substantially between ARG classes and was highest for the aminoglycoside resistance gene AAC(3) while the levels for beta-lactamases were, generally, lower. ARGs involved in IPTs also differed between phyla where, for example, tetracycline resistance genes were commonly transferred between Firmicutes and Proteobacteria, but rarely between Actinobacteria and Proteobacteria. The results, furthermore, show that conjugative systems are seldom shared between bacterial phyla, suggesting that other mechanisms drive the dissemination of ARGs between divergent hosts. We also show that bacterial genomes involved in IPTs of ARGs are either over- or under-represented in specific environments. These IPTs were also found to be more recent compared to transfers associated with bacteria isolated from water, soil, and sediment. While macrolide and tetracycline resistance genes involved in ITPs almost always were +95% identical between phyla, corresponding β-lactamases showed a median identity of < 60%. We conclude that inter-phyla transfer is recurrent and our results offer new insights into how resistance genes are disseminated between evolutionary distant bacteria.

## Introduction

Infections from antibiotic-resistant bacteria constitute a growing global public health crisis which in 2019 alone was associated with almost 5 million deaths [1]. Bacteria often become resistant to antibiotics by acquiring antibiotic resistance genes (ARGs) through the process of horizontal gene transfer [2]. ARGs are commonly located on mobile genetic elements (MGEs), such as plasmids and other integrative elements, transposons, and integrons, which allow them to efficiently move within and between bacterial cells [3]. Many ARGs, encoding a wide range of resistance mechanisms, have been described to date [4]. This number is constantly increasing, not only due to the discovery of ARGs in pathogenic bacteria but also because new resistance genes are frequently identified in non-pathogenic bacteria and in metagenomes, where it is often not possible to assign a specific host [5,6].

ARGs are commonly shared between evolutionary distant hosts. For example, the New Delhi metallo-β-lactamase (NDM), which provides resistance to penicillin, cephalosporins, and carbapenems, has been detected in multiple bacterial phyla, including pathogens from Proteobacteria, Bacteroidetes, and Firmicutes [7,8]. Similarly, the monooxygenase *tet(X)*, which provides high-level resistance by degrading both tetracyclines and tigecycline, is hypothesized to originate from Flavobacteriaceae (Bacteroidetes) but, despite this, *tet(X)* is commonly found in species from both Proteobacteria and Firmicutes [9]. A third example is the macrolide resistance gene *erm(B)*, which is well-spread in Proteobacteria but has been suggested to originate from a yet undiscovered Firmicutes species [10]. The recruitment of ARGs from evolutionary distant bacteria has, thus, enabled pathogens and other bacteria to adapt to strong antibiotic selection pressures and constitutes a major component in the development of multiresistant strains.

The origins of most ARGs encountered to date remain unknown. Indeed, less than 5% of the ARGs have been associated with a host from which it was mobilized onto MGEs that later spread horizontally to pathogens [11]. Bacterial communities, both in the human and animal microbiome and in external environments, such as soil, water, and sediments, are known to harbor large and diverse collections of ARGs, many of today remain uncharacterized [12]. These ARGs have been hypothesized to constitute a reservoir from which novel resistance determinants can be recruited and, eventually, transferred into pathogens [13]. This will, in many cases, require transfers between evolutionary distant bacterial hosts – a process that, to a large extent, remains unknown.

In this study, we aim to systematically analyze the transfer of ARGs between bacterial phyla. We also aimed to describe taxonomical patterns of the inter-phyla flow of ARGs and to identify environments where these transfers are most likely to occur. We took advantage of the large number of sequenced genomes currently present in public repositories [5,14,15] and the development of accurate computational methods to identify both established and previously uncharacterized resistance determinants [5,16], to analyze almost half a million bacterial genomes for the horizontal transfer of ARGs. Through analysis of inconsistencies between host taxonomy in phylogenetic trees constructed from ARGs, we could identify 661 inter-phyla transfers (IPTs) of ARGs. Our results show that most types of ARGs have been involved in IPTs, but the observed frequency of IPTs varied between resistance mechanisms. Network analysis revealed that Proteobacteria was the most connected phylum, followed by Firmicutes and Actinobacteria. By contrast, low frequencies were observed for IPTs involving Bacteroidetes, especially for those also involving Actinobacteria. Finally, our results suggest that IPTs were associated with different environments depending on the resistance mechanism, implying that ecological and taxonomic factors play important roles.

## Results

### Antibiotic resistance genes are ubiquitously present in all major bacterial phyla

A total of 427,495 bacterial genomes were screened for antibiotic resistance genes (ARGs) using fARGene, a software that uses optimized probabilistic models to identify resistance genes in sequence data [16]. Each genome was analyzed for 22 classes of resistance genes, representing 18 resistance mechanisms against five major groups of clinically relevant antibiotics (aminoglycosides, β-lactams, fluoroquinolones, macrolides, and tetracyclines), resulting in almost one million (993,827) significant matches. The number of identified ARGs differed between gene classes, where AAC(6’) aminoglycoside acetyltransferases were the most common (180,321 matches), while tetracycline degradation enzymes were the least common (1,049 matches). ARGs were found in genomes from all the largest phyla, most commonly in Proteobacteria (66.3% on average) followed by Firmicutes (20.4%), Actinobacteria (8.4%), Bacteroidetes (3.9%), Cyanobacteria (0.22%), and Chloroflexi (0.18%) (Fig 1A).

**Fig 1.**
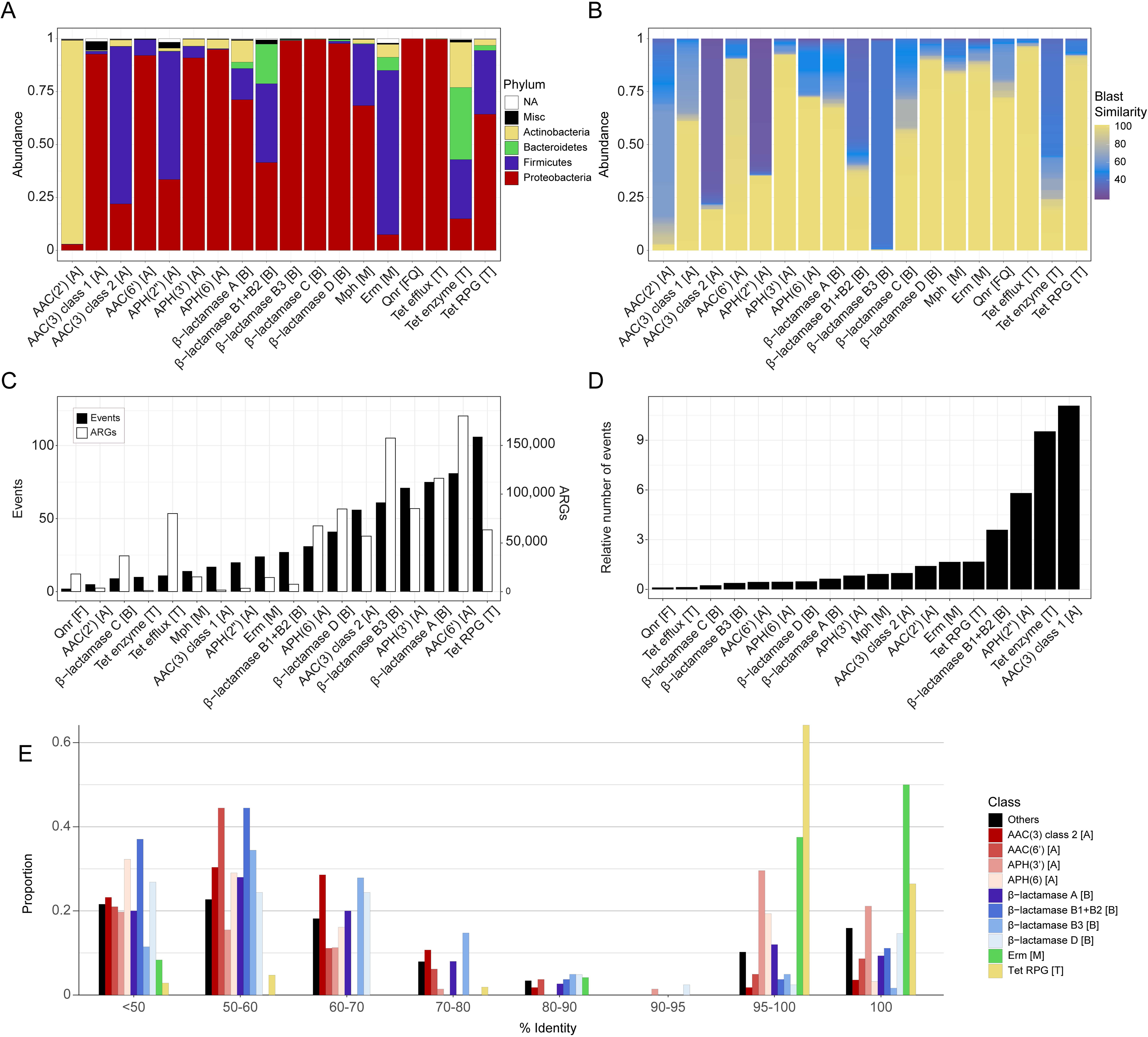
Overview of the identified antibiotic resistance genes (ARGs) and detected inter-phyla transfers (IPTs). The distribution of host phyla (A) differed considerably between ARGs. In (B), the sequence similarity of the identified ARGs when compared to genes present in the ResFinder database is shown. The number of IPTs, together with the number of detected ARGs is shown in (C) while the ratio between the number of IPTs and the number of detected ARGs within each group, multiplied by 1000, is shown in (D). Finally, in (E) the maximum sequence similarity of ARGs involved in IPTs from different phyla is shown, suggesting both recent and more ancient transfers. The letters in the brackets after the gene names indicate the class of antibiotics for which they provide resistance: A for aminoglycosides, B for beta-lactams, M for macrolides, and T for tetracyclines.

The phyla of the hosts carrying ARGs varied between gene classes. Most ARGs were predominantly found in Proteobacteria but there were exceptions, such as AAC(2’) which was almost exclusively found in Actinobacteria (96.2% of the AAC(2’)-encoding species) and AAC(3) (of class 2), APH(2”), and Erm which were mostly found in Firmicutes (74.4%, 60.5% and 77.6% of the species, respectively). Bacteroidetes, in which ARGs were less commonly observed, had a high prevalence of genes encoding tetracycline degradation enzymes and class B1/B2 β−lactamases (present in 34.0% and 18.7% of the species, respectively). Furthermore, most of the identified ARGs were identical or highly similar to previously well-characterized resistance genes (Fig 1B). There were, however, a substantial proportion of less well-described genes – especially encoding class B1/B2 and B3 β−lactamases, tetracycline degradation enzymes, and aminoglycoside modifying enzymes AAC(2’), APH(2’’), and AAC(3) which is in line with previous studies [10,17,18]. In total, close to two-thirds (66.2%) of the ARGs showed a high sequence similarity (≥ 90%) to at least one well-characterized ARG.

### Detection of inter-phyla transfer of ARGs

We implemented and applied an algorithm to identify inter-phyla transfers (IPTs) of ARGs. The algorithm was based on gene-specific phylogenetic trees (S1-18 Figs), in which IPTs were detected based on differences in the taxonomy of the bacterial hosts carrying evolutionary similar ARGs (see Methods). As a part of this process, we re-evaluated the taxonomic affiliation of the bacterial genomes appearing in sparsely populated leaves of the ARG trees to minimize the impact of false positives (see S1 Table for the 1,221 excluded genomes).

In total, we detected 661 IPTs of ARGs, which were unevenly distributed between the resistance gene classes (Fig 1C). The highest number of IPTs was found for the tetracycline ribosomal protection genes (RPGs; 106 transfers), followed by aminoglycoside acetyltransferase AAC(6’) (81 transfers), and class A beta-lactamases (75 transfers), while, in contrast, only 2 transfers were detected for quinolone resistance (*qnr*) genes. There was, as expected, a positive correlation between the number of observed IPTs and the number of identified ARGs (r=0.71, p=6.3×10^-4^), however, this trend disappeared after normalization by the ARG frequency (Fig 1D). The aminoglycoside acetyltransferases AAC(3) (type 1) and APH(2’’), along with the tetracycline-degrading enzymes, had the highest number of IPTs in relation to the identified ARGs. Although these genes were relatively infrequent, with 1,534, 3,441, and 1,049 identified genes, respectively, they were associated with a substantial portion of the total IPTs, corresponding to 17, 20, and 10 IPTEs, respectively (Fig 1D). Interestingly, all classes of β-lactamases, except for B1/B2, were associated with a relatively low number of IPTs compared to their prevalence (Fig 1D).

The sequence similarity of ARGs involved in IPTs in their different phyla varied (Fig 1E). Most RPG, Erm, and APH(3’) genes were found to be similar between phyla (median amino acid similarity 99.7%, 99.8%, and 98.16% respectively), suggesting that they were more recently transferred. In contrast, β-lactamases and other aminoglycoside-modifying enzymes showed, in general, a larger difference (median sequence similarity between 59.1% and 57.3%).

### The structure of the inter-phyla transfers of ARGs

Network analysis was used to visualize how ARGs have been transferred between phyla (Fig 2). This showed that Proteobacteria had the largest number of connections, primarily to Firmicutes, Actinobacteria, and Acidobacteria (136, 56, and 48 IPTs, respectively). Transfers between Proteobacteria and Chloroflexi, Cyanobacteria, and Verrucomicrobia were also observed, but less frequently (11, 11, and 26 IPTs, respectively). Stratification of the results into resistance gene classes showed that Proteobacteria played a central role for all ARGs except for Erm and RPG (Fig 2C–F). Aside from Proteobacteria, IPTs between Firmicutes and Actinobacteria (49) and between Firmicutes and Bacteroidetes (21) were also commonly observed. Interestingly, only a single IPT was detected between Actinobacteria and Bacteroidetes.

**Fig 2.**
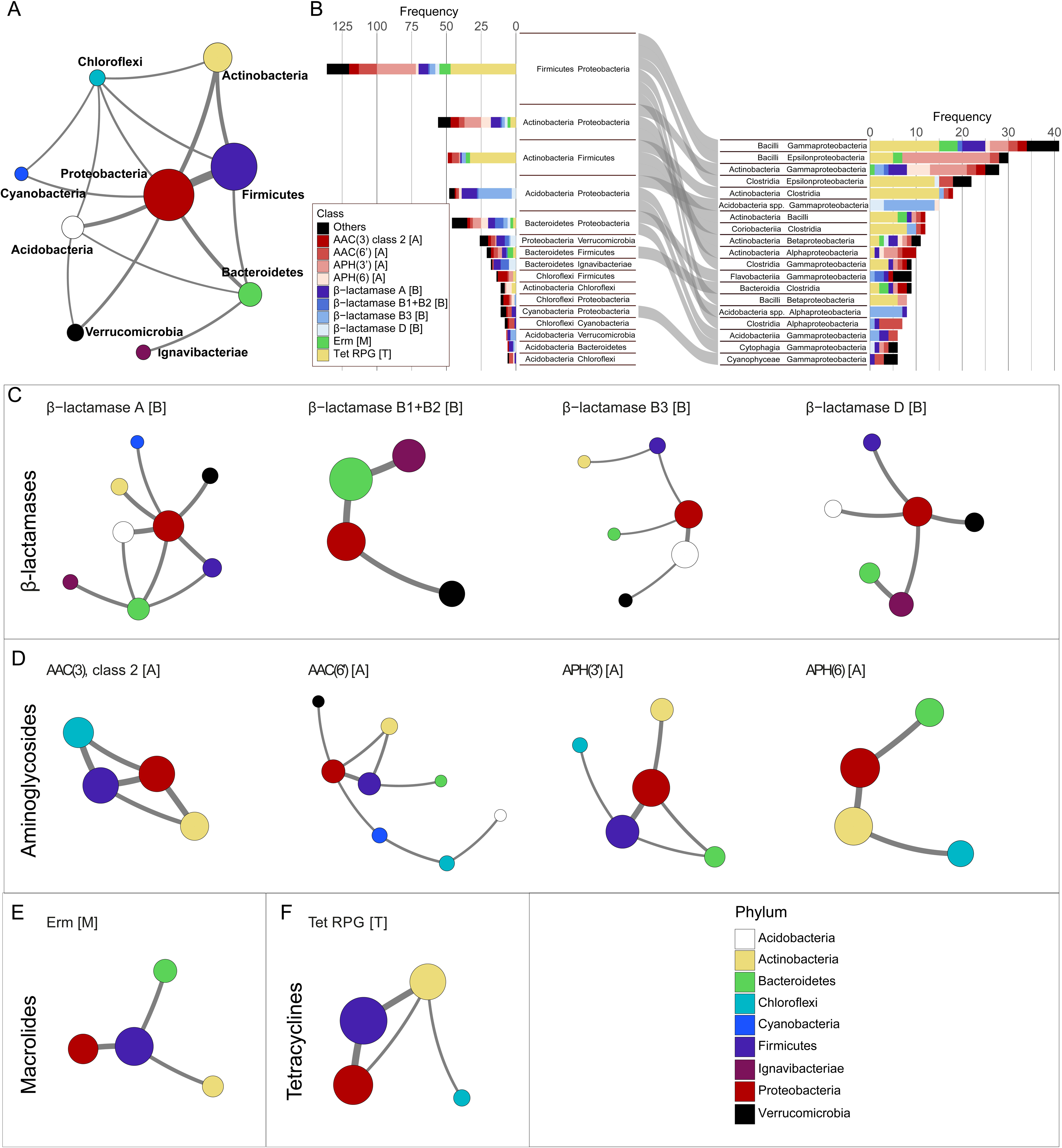
Analysis of the transfer of ARGs. In (A), a network representation of the interphyla transfers (IPTs) between Proteobacteria, Firmicutes, Actinobacteria, Chloroflexi, Cyanobacteria, Acidobacteria, Verrucomicrobia, Bacteroidetes, and Firmicutes. In (B), the most common IPTs are stratified based on the involved ARGs. Panels (C) to (F) show gene-specific transfer networks. Ribosomal protection genes are abbreviated RPG. The letters in the brackets after the gene names indicate the class of antibiotics for which they provide resistance: A for aminoglycosides, B for beta-lactams, M for macrolides, and T for tetracyclines.

The transfer of β-lactamases was predominantly observed between Proteobacteria and other Gram-negative phyla, especially Bacteroidetes, Acidobacteria, and Verrucomicrobia (Fig 2C). We noted that the relatively rare class B1+B2 β-lactamases have been commonly transferred between Proteobacteria and Bacteroidetes (6 IPTs), more than the frequently encountered class A β-lactamases (5 IPTs). Among the aminoglycoside resistance genes, AAC(3), AAC(6’), and APH(3’) were all observed to be transferred between Firmicutes and Proteobacteria (7, 13, and 28 IPTs, respectively), while no such transfers were detected for APH(6). Transfers involving Bacteroidetes could be seen for all aminoglycoside resistance mechanisms except AAC(3’) where instead transfers involving Chloroflexi were frequent (Fig 2D). The transfers of the Erm macrolide resistance mechanism involved, almost exclusively, Firmicutes and either Proteobacteria, Bacteroidetes, or Actinobacteria (Fig 2E). Similarly, tetracycline RPGs were frequently transferred between Firmicutes and Proteobacteria and between Actinobacteria and Firmicutes, but rarely between Firmicutes and Bacteroidetes (Fig 2F). Finally, we noted that Erm and RPGs had either very few or no detected transfers between Proteobacteria and Actinobacteria, even though these classes of ARGs were common in both phyla.

Analysis of the IPTs at higher taxonomic resolution revealed distinct patterns (Fig 2B). The class Bacilli (Firmicutes) was involved in the highest number of transfers, primarily together with Gamma- and Epsilon-proteobacteria. Interestingly, a large proportion of the transfers between Bacilli and Gammaproteobacteria was associated with tetracycline RPGs while the transfers between Bacilli and Epsilonproteobacteria were instead dominated by aminoglycoside modifying enzymes. The transfers of RPGs to and from Epsilonproteobacteria were, instead, frequently involving Clostridia (Firmicutes). The IPTs including Proteobacteria and Actinobacteria were primarily observed between Alpha-, Beta- and/or Gammaproteobacteria and the eponymous Actinobacteria class. We noted, however, that not a single transfer of RPGs could be detected between Gammaproteobacteria and Actinobacteria (class) while, in contrast, these genes were commonly transferred between Actinobacteria (class) and Clostridia. The transfer between Coriobacteria (Actinobacteria) and Clostridia was also dominated by RPGs, but not a single IPT could be detected between Coriobacteria and any proteobacterial class.

### Conjugative elements associated with the inter-phyla transfer of ARGs

Conjugative elements, in particular plasmids, have been hypothesized to play a central role in the inter-phyla transfer of bacterial genes [19,20]. We, therefore, annotated the genomic context of all ARGs associated with IPTs for genes encoding mating pair formation (MPF) proteins and relaxases, which are vital for pilus formation and DNA mobilization in conjugative transposition, respectively (Methods) [21]. Our results showed that the class of MPF proteins varied substantially between phyla, where all included types could be found in proteobacterial genomes while the classes FA and FATA were, almost exclusively, found in Firmicutes, Actinobacteria, and Bacteroidetes (Fig 3A). Relaxases were, generally, more spread, where four types (MOB_F_, MOB_P_, MOB_Q_, MOB_V_) were commonly found in two or more phyla.

**Fig 3.**
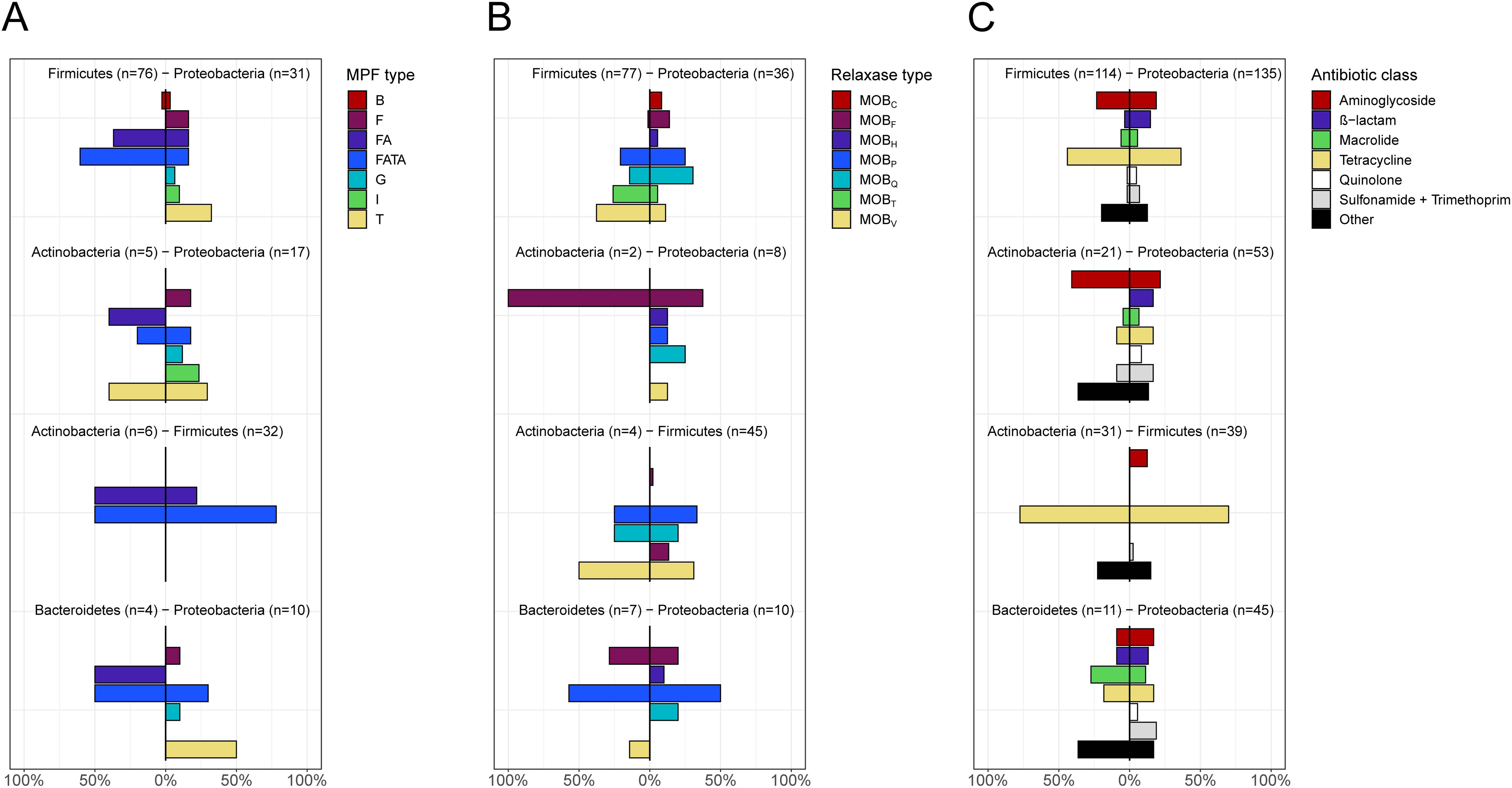
The distribution of genes involved in conjugation and co-localized resistance genes in the genetic context of ARGs involved in IPTs. The proportion of (A) MPF genes, (B) relaxases, and (C) co-localized resistance genes identified in the genetic context of the ARGs involved IPTs, for different pairs of phyla.

We, furthermore, investigated if ARGs transferred between two phyla were associated with the same type of conjugative elements in their respective host genomes (Methods). For MPF proteins, no clear general pattern could be identified (Fig 3A, p=0.071, permutation test), however, relaxases showed a significant similarity (Fig 3B, p=0.0068, permutation test) where the same type of MOB gene was often co-localized with the same ARG in hosts from both phyla. This could, for example, be seen for Actinobacteria and Firmicutes (MOB_P_, MOB_Q_, MOB_V_) and Proteobacteria and Bacteroidetes (MOB_P_) (S19 Fig) Even more distinct similarities could be seen for the co-localization of the IPT-associated ARGs with other resistance genes (Fig 3C, p=6.0×10^-4^, permutation test). Here, the co-localization of tetracycline resistance genes in IPT between Actinobacteria and Firmicutes was most prominent. These results suggest that IPTs may be mediated by non-conjugative plasmids carrying relaxases that are compatible over large evolutionary distances. It also suggests that transfers often include larger genomic regions, containing more than a single ARG.

### Inter-phyla transfers are overrepresented and recent in the human microbiome

Next, the isolation sources of the bacterial genomes carrying ARGs involved in IPT were examined to assess where these bacteria are especially common. The human microbiome was found to be the most frequent isolation source, from which 52.8% of the genomes carrying an ARG involved in IPT were isolated. This was followed by soil (19.4%), water (12.0%), animal (10.0%), and, finally, sediment (2.5%). Statistical analysis showed that IPTs of ARGs were significantly enriched with specific environments (Fig 4). In particular, Mph, class B1/B2 and D β-lactamases, and tetracycline RPGs were all found in sequenced genomes isolated from multiple environments while the genomes carrying the ARGs involved in IPTs were highly overrepresented in the human microbiome (p<10^-15^ for all cases, Fig 4B). In contrast, IPTs involving AAC(6’) and APH(6) aminoglycoside-modifying enzymes, class A β-lactamases, and tetracycline efflux pumps were underrepresented in the human microbiome (p<10^-15^ for all cases).

**Fig 4.**
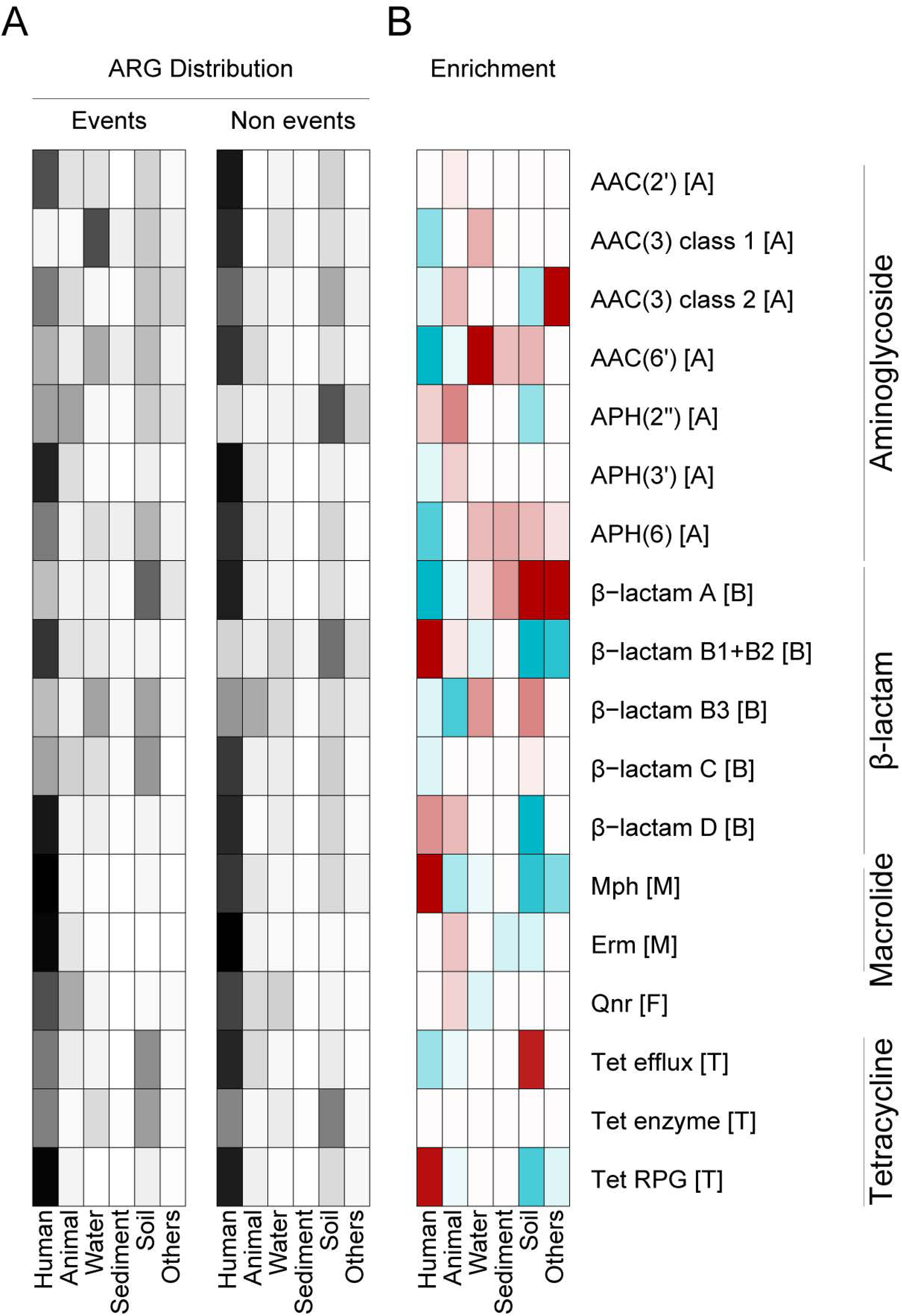
Heatmaps describing the distribution and enrichment of hosts carrying ARGs involved in IPT stratified based on isolation source and gene class. In (A), the distribution of hosts involved and not involved in IPTs are shown. In (B), the enrichment score describing the statistical overrepresentation (red) and underrepresentation (blue) of hosts involved in IPTs are shown. The overrepresentation was assessed by Fisher’s exact test. Tests with p>0.05 were marked as white and not considered to be significant. The letters in the brackets after the gene names indicate the class of antibiotics for which they provide resistance: A for aminoglycosides, B for beta-lactams, M for macrolides, and T for tetracyclines.

Several ARGs involved in IPT were also carried by bacterial genomes that were overrepresented in the external environments. For example, the hosts carrying AAC(6’) (p<10^-15^), APH(6) (p=1.6×10^-14^), and class B3 β-lactamases (p<10^-15^) involved in IPTs were overrepresentation in water while the IPT-involved hosts for class A β-lactamases (p<10^-15^) and tetracycline efflux pumps (p<10^-15^) and, to a less extent, also AAC(6’) and APH(6) (p<10^-15^ and p=4.7×10^-14^, respectively) were overrepresented in soil. Interestingly, the IPTs for two ARGs – AAC(3) (type 2) and class A β-lactamases – were significantly associated with hosts isolated from ‘food’ and ‘milk’ (p<10^-15^ for both gene classes).

Finally, we noted that the sequence similarity of ARGs involved in IPTs varied significantly between different phyla across environments (Fig 5). Most similar ARGs were found in the human and animal microbiome (median similarity of 99.22% and 98.98%, respectively), suggesting that these transfers are more recent. Compared to the human microbiome, the ARGs transfers in water, sediment, and soil were all significantly more different in their respective phyla, which may suggest domination by more ancient transfers (median similarity 56.33% (p = 2.41e-10), 53.03% (p < 1.76e-08), 53.42% (p < 6.93e-14), respectively, Wilcoxon’s rank sum test).

**Fig 5.**
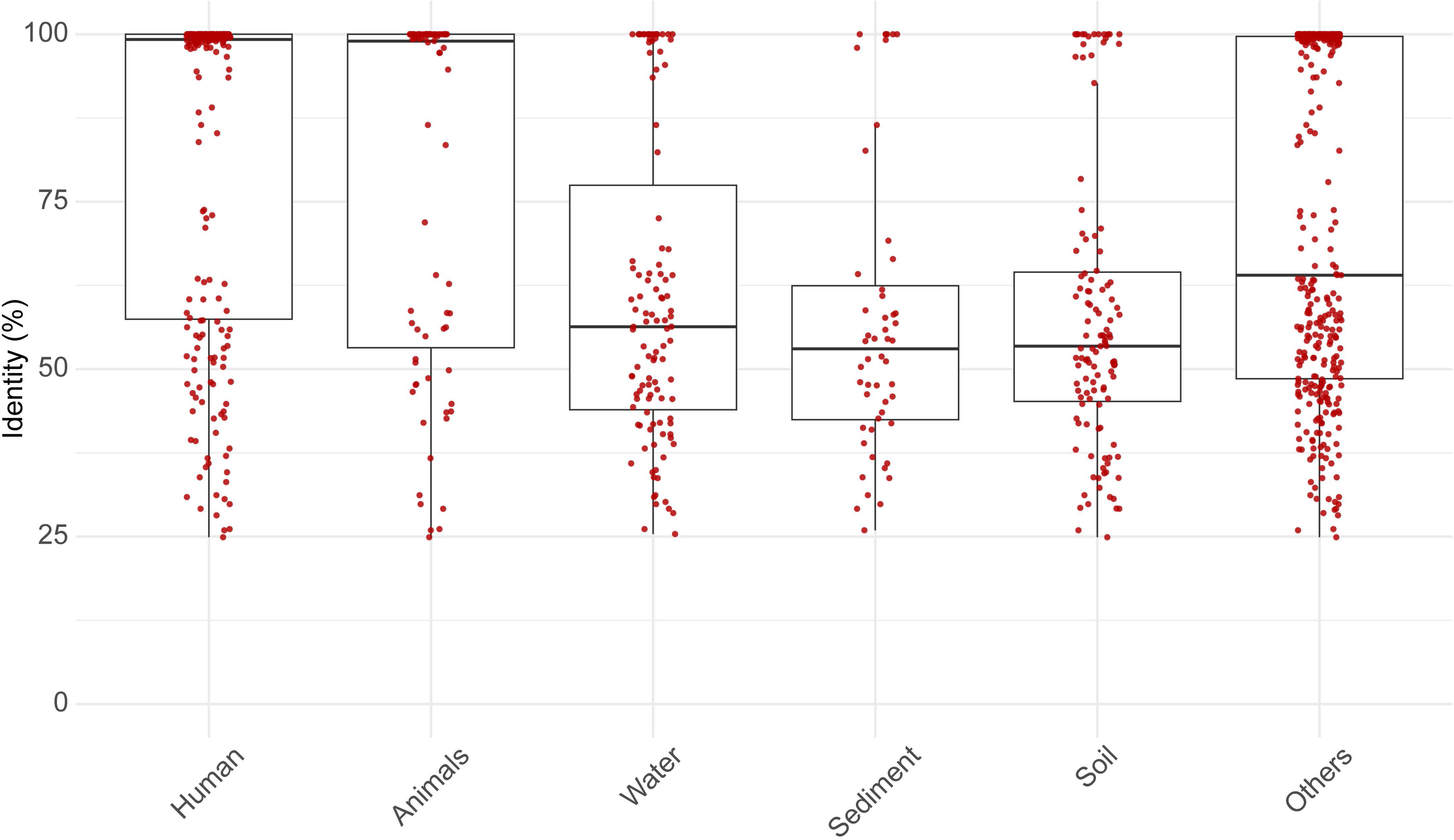
ARGs involved in inter-phyla transfers are more similar between their different phyla in the human and animal microbiome compared to the environment. Genomes were classified into environments based on their isolation source: human microbiome, animal microbiome, water, sediment, soil, and, others. The boxplots represent the distribution of the sequence identity across each environment. The individual data points, shown as red jittered dots, illustrate the distribution within each group.

## Discussion

Antibiotic resistance genes (ARGs) are transferred from distantly related bacteria into pathogens, which makes them harder to treat. Inter-phyla transfers (IPTs) of ARGs have been repeatedly documented in the literature [22–24] but the knowledge of which gene, bacteria, and environments are involved in these gene flows has so far been limited. In this study, we systematically investigated ARGs that have undergone horizontal transfer between phyla, representing the highest taxonomic level within the bacterial domain. The analysis, which was based on almost one million resistance genes identified in more than 400,000 bacterial genomes, showed that the majority of the 22 analyzed ARG classes were subjected to IPTs. The results also showed that the frequency of IPTs varied substantially between antibiotic and ARG classes, with high frequencies observed for aminoglycoside, tetracycline, and macrolide resistance genes. Here, AAC(3) showed the highest relative frequency, where one out of every hundred encountered genes were associated with an IPT. All classes of ARGs were, furthermore, associated with recent events where the gene sequences present in both involved phyla were identical, or close to identical. However, for many resistance mechanisms, especially β-lactamases, we could also identify evolutionary older IPTs, suggesting that inter-phyla transfer of ARGs is ancient and, thus, not only a consequence of the antibiotic mass consumption during the last hundred years.

All major bacterial phyla were involved in the inter-phyla transfer of ARGs. An especially large number of gene transfers were associated with Proteobacteria, Firmicutes, and Actinobacteria, which are highly abundant in the genome databases, but IPTs were also detected for less frequently sequenced bacteria, such as Bacteroidetes, Chloroflexi, Cyanobacteria, and Verrucomicrobia. For several of the analyzed ARGs, Proteobacteria acted as a central hub with connections to multiple phyla. These connections were, to a large extent, associated with bacterial hosts from human and animal microbiomes, but also include genomes isolated from the external environment, such as soil and water (S20-37 Figs). These results, thus, reaffirm the plasticity of many proteobacterial genomes and show their ability to share antibiotic-resistance genes over large evolutionary distances. Indeed, pathogens from Proteobacteria, such as *Escherichia coli*, *Klebsiella pneumoniae, and Pseudomonas aeruginosa*, commonly carry ARGs that are hypothesized to originate from other phyla (e.g. and *erm(B)*) [25–27]. Proteobacteria harbor broad host range conjugative elements, some of which are known to be able to move across large evolutionary distances. This includes, for example, plasmids carrying class T mate-pair forming (MPF) genes, which have previously been documented also in e.g. Actinobacteria [20]. We found that class T MPFs were commonly co-localized with ARGs associated with IPT between Proteobacteria and Actinobacteria, indicating that it may be one of the mechanisms that mediate gene transfers between these phyla. In contrast, there are currently no documented conjugative plasmids that are commonly present in both Proteobacteria and Firmicutes [20]. This is also in line with our findings, where plasmids carrying FA and FATA MPF, which are common in Firmicutes, were rare in Protobacterial hosts. Despite this, we detected a large gene flow between Proteobacteria and Firmicutes, suggesting that other mechanisms may be used for these transfers. We noted, however, that the genetic context of ARGs associated with IPTs displayed similarities, frequently showing co-localization of relaxases and other resistance genes across different phyla. This suggests that the horizontal transfer between phyla, including those between Proteobacteria and Firmicutes, likely includes larger genetic regions, and, potentially, involves non-conjugative plasmids.

Bacteroidetes were involved in a low number of inter-phyla transfers. The few IPTs involving Bacteroidetes included, in addition to the β-lactamases transferred to and/or from Proteobacteria, a diverse set of ARGs that were also shared with Firmicutes. Interestingly, this included genes conferring resistance to aminoglycosides (e.g. APH(3’) and APH(6)), a class of antibiotics for which many Bacteroidetes are known to be intrinsically resistant [28]. These transfers may result from co-selection, potentially through co-localization of aminoglycoside resistance genes and other ARGs on the same MGE. Another possibility is that aminoglycoside resistance genes still provide a significant fitness advantage in Bacteroidetes – a hypothesis that is supported by a recent study that showed that aminoglycoside resistance genes from Bacteroidetes can provide clinical levels of resistance in *Escherichia coli* [10]. We, furthermore, only observed two IPTs between Bacteroidetes and Actinobacteria, even though bacteria from these phyla are common in both host-associated and external environments [29,30]. Indeed, Bacteroidetes (e.g. *Bacteroides spp.*) and Actinobacteria (e.g. *Bifidobacterium spp.*) are integral parts of the human gut microbiome, suggesting that they are commonly co-occurring in environments that recurrently are under strong selection pressures for antibiotic resistance [31]. The lack of detected transfers suggests that there are barriers that limit the exchange of ARGs between these phyla. We found that Actinobacteria and Bacteroidetes both carried conjugative elements with class FA and FATA MFP, however, previous studies have shown that are generally rare on plasmids in Bacteroidetes [20]. Genomes from Bacteroidetes are, furthermore, typically AT-rich while Actinobacteria are typically GC-rich, suggesting that low gene compatibility may make it hard to find ARGs that have the necessary efficiency in both phyla [4]. Indeed, many ARGs depend on high expression to induce a sufficiently strong resistance phenotype and an inefficient codon configuration may, thus, result in too high fitness costs [32,33]. Nevertheless, Bacteroidetes have been estimated to have a substantially higher rate of horizontal gene transfer than other common members of the human gut microflora [29], still, our results suggest that this does not include inter-phyla transfers of antibiotic resistance genes.

Strong selection pressures likely govern the flow of ARGs between phyla. This was seen for all classes of β-lactamases, which are enzymes that break down β-lactams, and the primary means of resistance against these antibiotics in Gram-negatives [34]. Our results showed that β-lactamases were almost exclusively transferred between Proteobacteria, Acidobacteria, Bacteroidetes, and Verrucomicrobia, all of which predominantly include Gram-negative bacteria. A similar pattern could be seen for aminoglycoside resistance transferred between Proteobacteria and the two Firmicute classes Bacilli and Clostridia. Aminoglycosides use active electron transports to enter the cell [35] and are therefore highly effective against aerobes, such as Bacilli, while the potency against anaerobes, such as Clostridia [36], is typically low. This was reflected in the gene flow, where the transfer of aminoglycoside resistance genes (especially APH(3’)) was observed between Proteobacteria and Bacilli, but not between Proteobacteria and Clostridia. Our results also suggest that IPTs of some classes of ARGs may be associated with specific environments. For example, the macrolide resistance enzyme Mph was detected in both the human microbiome and the external environment. A similar pattern was seen for tetracycline RPG, which was associated with IPTs involving hosts that were highly over-represented in the human microbiome. Interestingly, our results also indicated that the ARGs involved in IPT were more similar between phyla in the human and animal microbiome compared to the external environment. Strong selection pressures have been shown to promote horizontal gene transfer [37], suggesting that the IPTs seen between bacteria present in the host-associated bacterial communities may be a consequence of the last 80 years of mass consumption of antibiotics.

We, finally, noticed that the transfer of RPGs was particularly common between Proteobacteria and Firmicutes (especially between Clostridia and Epsilonproteobacteria), Actinobacteria and Firmicutes but, interestingly, very rare between Proteobacteria and Actinobacteria (Fig 2A). Successful inter-phyla transfer requires that the involved hosts are physically present in the same bacterial community. It is, in this context, worth noting that Epsilonproteobacteria and Clostridia are both common in the microbiome of poultry [38,39], for which a significant proportion of the produced tetracyclines is used for growth promotion [40]. We could, however, not statistically assess overrepresentation for this particular environment due to relatively few bacterial isolates with a specified isolation source.

In this study, we use phylogenetic trees reconstructed from half a million ARG sequences to detect horizontal transfers between bacterial phyla. In contrast to many previous studies of horizontal gene transfer, our method is not dependent on the identification of known MGEs and is thus more general. Indeed, many of the MGEs associated with horizontal transfer of ARGs can excise themselves from the genome, leaving no or very few changes in the nucleotide sequence of the host. Many MGEs are still uncharacterized and thus not properly annotated in existing sequencing databases. This often makes the association between horizontal gene transfers and MGEs difficult to establish [2]. It should, however, be noted that our method is dependent on the correct taxonomic affiliation of the included genomes. Since erroneously annotated sequences and genomes are common in GenBank, we applied three independent methods (alignment to the SILVA 16S database, Metaxa2, GTDBK) to scrutinize the taxonomic affiliation [41–43]. To minimize the proportion of incorrectly annotated genomes – and thus the number of falsely predicted IPTs – we excluded all genomes for which the phylum was uncertain. Moreover, our results are highly dependent on the content in the genome databases. Indeed, we can, only report transfers that are documented, and, thus, the number of IPTs is thus likely underestimated for many parts of the taxonomic tree. Throughout the paper, we therefore represent the number of IPTs in relative terms, either compared to the total number of ARGs or, as in Fig 5, in relation to the genomes not associated with IPTs. Considering that current databases only reflect a small part of the total microbial diversity on Earth, our estimates, therefore, should be considered conservative and revisited as more genomes become available in the sequence repositories.

ARGs are often transferred from evolutionarily distant species before they are acquired by pathogens. We know, however, little about this process, especially regarding the transfers between the most divergent bacterial hosts. In this study, we provide a comprehensive and systematic analysis of the inter-phyla transfer of ARGs reflecting the body of knowledge currently available in sequence repositories. We demonstrate that inter-phyla transfer is a widespread phenomenon in most parts of the bacterial tree of life and encompasses multiple clinically relevant resistance mechanisms. Recent inter-phyla transfers were, furthermore, found to be especially common in the human microbiome, likely promoted by decades of antibiotic mass consumption. Our study, thus, provides new insights into the evolutionary processes that result in multiresistant pathogens through the accumulation of resistance genes. We conclude that the development of management strategies that efficiently limit the spread of ARGs is vital to ensure the potency of both existing and future antibiotics.

## Material and Methods

### Prediction of antibiotic resistance genes in bacterial genomes

A total of 427,495 bacterial genomes encompassing 47,582,748 sequences were downloaded from NCBI GenBank (October 2019) [44] and analyzed using fARGene (v0.1, default parameters) [45]. fARGene was executed using 22 profile hidden Markov models (HMMs) [45,46] built to identify genes encoding 18 resistance mechanisms against five major classes of antibiotics: β-lactamases (class A, B1+B2, B3, C, and D); macrolides (Erm 23S rRNA methyltransferases and Mph macrolide 2’-phosphotransferases); tetracyclines (efflux pumps, ribosomal protection genes (RPGs), and drug-inactivating enzymes); fluoroquinolones (Qnr) and aminoglycosides (aminoglycoside-modifying enzymes including AAC(2’), AAC(3)-class 1, AAC(3)-class 2, AAC(6’), APH(2”), APH(3’), and APH(6)). All matches above the default profile-specific significance threshold were putative ARGs and stored for further analysis.

### Phylogenetic analysis and prediction of inter-phyla transfers

The ARGs predicted by each gene model were aligned using Clustal Omega (v1.2.4, default parameters) [46]. Each alignment was then used to calculate an unrooted phylogenetic tree using the maximum-likelihood algorithm implemented in FastTree (v2.1.10, all other parameters set to default) [47]. The taxonomy for each sequence was retrieved from the NCBI taxonomy database using “accessionToTaxa” and “getTaxonomy” from the R package “taxonomizr” (v0.5.3) [48].

Inter-phyla transfers (IPTs) were detected through a custom-built algorithm by identifying node points within the tree where the descendant hosts belong to at least two different phyla. Node points that contained an IPT in one of its subtrees were removed to avoid including the same leaves in multiple transfers and to keep only the evolutionarily most recent IPT for each gene. The algorithm was implemented in a custom-made script (https://git.io/JmVTa). The total list of host genome sequences descending from each node was obtained using the “Descendant” function from the “phangorn” R package (v2.5.5) [49]. Nodes were labeled from root to leaves using the “makeNodeLabel” function from the “ape” R package (v5.3) [50]. IPTs were visualized in phylogenetic trees plotted using the “ggtree” and “gheatmap” functions from the “ggtree” R package (v1.16.6) [51].

The taxonomic information reported in NCBI GenBank is known to occasionally suffer from misannotations. We, therefore, ensured that our analysis would be robust by scrutinizing the host taxonomy for IPTs where one or both phyla were represented by a limited number of genomes and where, thus, false positives may have a large effect (three or fewer genomes). All contigs from the genomes involved in these IPTEs were aligned to the SILVA 16S database (v138) [41] and the best hit was kept in each case, with the criteria of a maximum e-value of < 10^-3^ and a minimum coverage of 1,000 bp. In addition, we analyzed the sequences using Metaxa2 (v2.2) [42] (--mode genome) and with GTDBK (release 89) [52] (default parameters). Finally, all genomes annotated as “candidatus” were excluded from analysis but kept in the alignment and tree construction. By default, the NCBI taxonomy information was assigned to all contigs in an assembly project. If any of the SILVA, Metaxa2, or GRDBK results indicated a taxonomy different from NCBI, the assembly was removed from the remaining analysis. The genomes excluded due to questionable annotations can be found in the Supporting S1 Table.

For each predicted ARG sequence, the sequence similarity to well-characterized ARGs was calculated using “tblastn” from BLAST (v2.2.31+) [53], using ResFinder (downloaded 01-Oct-2019) as the reference database [15]. IPT networks were created based on the number of IPTs observed between different phylum pairs, either as an aggregate for all ARGs or as individual networks for each of the studied gene classes. Networks were plotted using the MapEquation InfoMap software (v.1.1.2) [54] with “--ftree” and “--no-infomap” parameters. To increase the readability, the networks were pruned. In the general network (Fig 2a), only edges corresponding to six or more observed IPTs were included. Similarly, in the gene class-specific networks (Fig 2B-F), only edges with three or more observed IPTs were included. The number of predicted ARGs from each mechanism were visualized as barplots for the taxonomic levels phylum and class using the R packages “ggplot2” (v3.3.2) [55] and “ggalluvial” (v0.12.3) [56]. In parallel, genetic distances between species from both branches of a certain node were compared at the amino acid level using “blastp” from BLAST (v2.2.31+) [53]. The maximum percent sequence identity for each node was saved and visualized as a histogram.

### Retrieval of isolation source information

For each identified host genome, information about its isolation source was retrieved from the NCBI nucleotide summary using the “esearch” function in the Entrez software (Version 11.8 (23 July 2019)) [57]. The reported isolation sources were manually scrutinized and categorized into five major groups: soil, sediment, water, human, animal, and others (S2 Table). Since some terms could be assigned into different categories and we wanted to preserve the individuality of the terms among the groups, we created an exclusion list to avoid misclassification (S3 Table).

Fisher’s exact test was used to assess the overrepresentation of genomes in their isolation source. Results were mapped using the “heatmap.2” function from the “gplots” (v3.6.3) [58] R package. Only results with a p-value < 0.05 were included in the final heatmap. Differences in ARG similarity between phyla across environments were assessed using the Wilcoxon rank-sum test.

### Genetic context analysis

Genetic context analysis was used to assess the mobility of the transferred ARGs. For each ARG included in an IPT, a region of up to 10kb up and downstream of the gene was retrieved from its host genome using GEnView v0.2 [59]. After retrieval, the genetic contexts were screened for the presence of genes associated with mobile genetic elements (MGEs). Genes involved in plasmid conjugation were identified by translating the genetic contexts in all six reading frames using EMBOSS Transeq v6.5.7.0 [60] and analyzing the translated sequences with 124 HMMs from MacSyfinder Conjscan v2.0 [61], using HMMER v3.1b2 [62]. Co-localized mobile ARGs were identified by finding the best among overlapping hits produced by “blastx” from BLAST (v2.10.1) [63], using a reference database of well-characterized ARGs based on ResFinder v4.0 [15], with the alignment criteria that hits should display >90% amino acid identity to a gene in ResFinder. The similarity of the distributions of MGEs and co-localized ARGs between phyla was tested using a permutation test. A similarity score was derived by adding the individual scores for each combination of phyla, which were calculated using Pearson’s χ^2^ test statistic. Phyla with very few (<5) observations (MGEs or ARGs) were excluded when calculating the similarity score to ensure a high statistical power. Significances were assessed from a null distribution derived by randomly permuting genomes within the same phyla 10,000 times.

## Supporting information

**S1 Table 1. Total number of removed sequences of taxonomy information for each class after following our curation criteria.** Sequences were kept in the alignment but did not compute for the evaluation of events.

**S2 Table. Isolation sources scrutinized from NCBI nucleotide database categorized into five major groups:** soil, sediment, water, human, animal, and others.

**S3 Table. Isolation source exclusion criteria.**

**S1–18 Figs. Circular phylogenetic trees representing each class of antibiotic resistance genes (ARGs).**

**S19 Fig.**

**S20-37 Figs. Association between each antibiotic resistance gene class and environments.**

