## Supplementary material for "The transfer of antibiotic resistance genes between evolutionary distant bacteria": S1-18 Figs

### aac2p

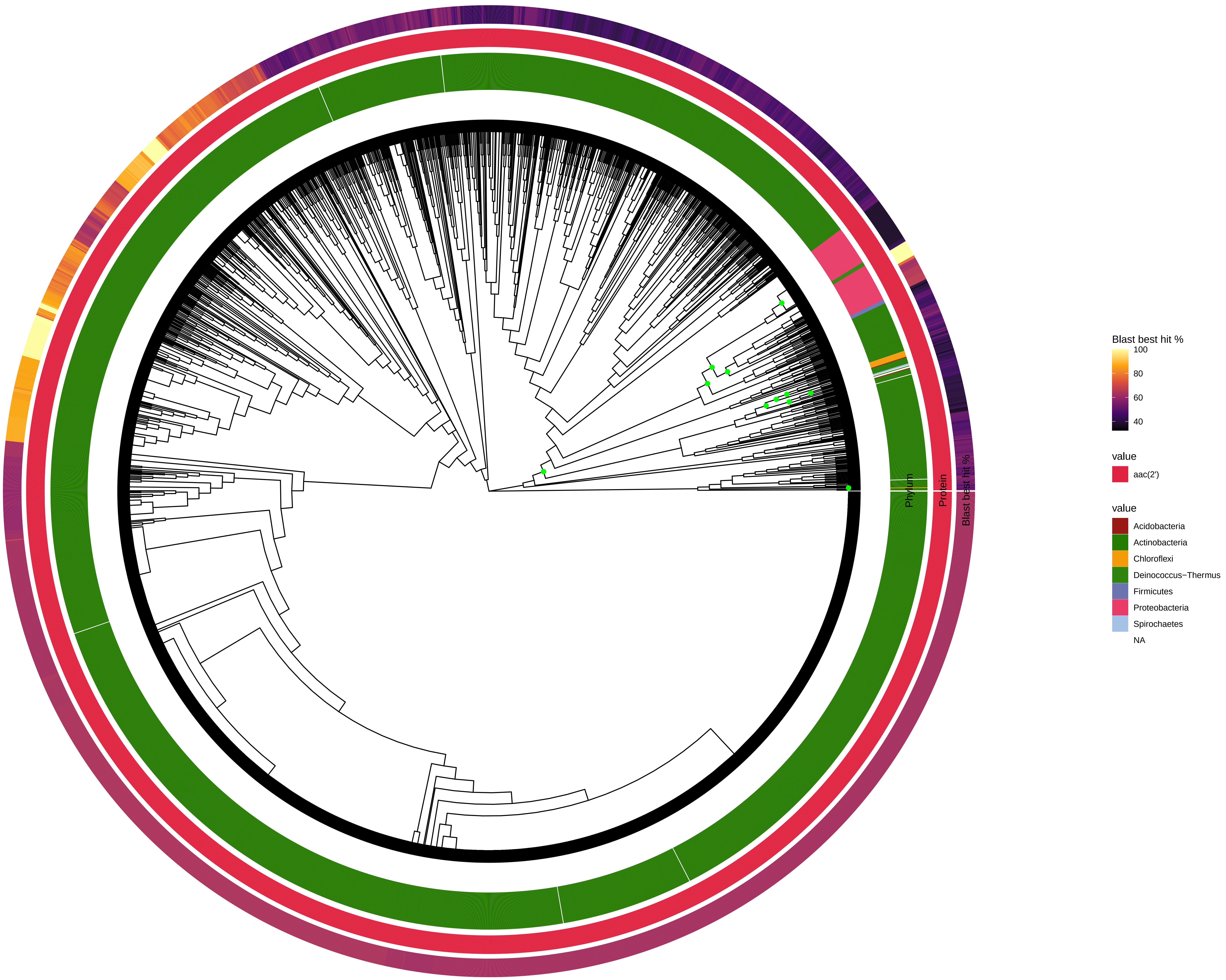

Phylum
Protein
Blast best hit %
Blast best hit %
100
80
60
40
value
aac(2')
value
Acidobacteria
Actinobacteria
Chloroflexi
Deinococcus−Thermus
Firmicutes
Proteobacteria
Spirochaetes
NA

### aac3_class1

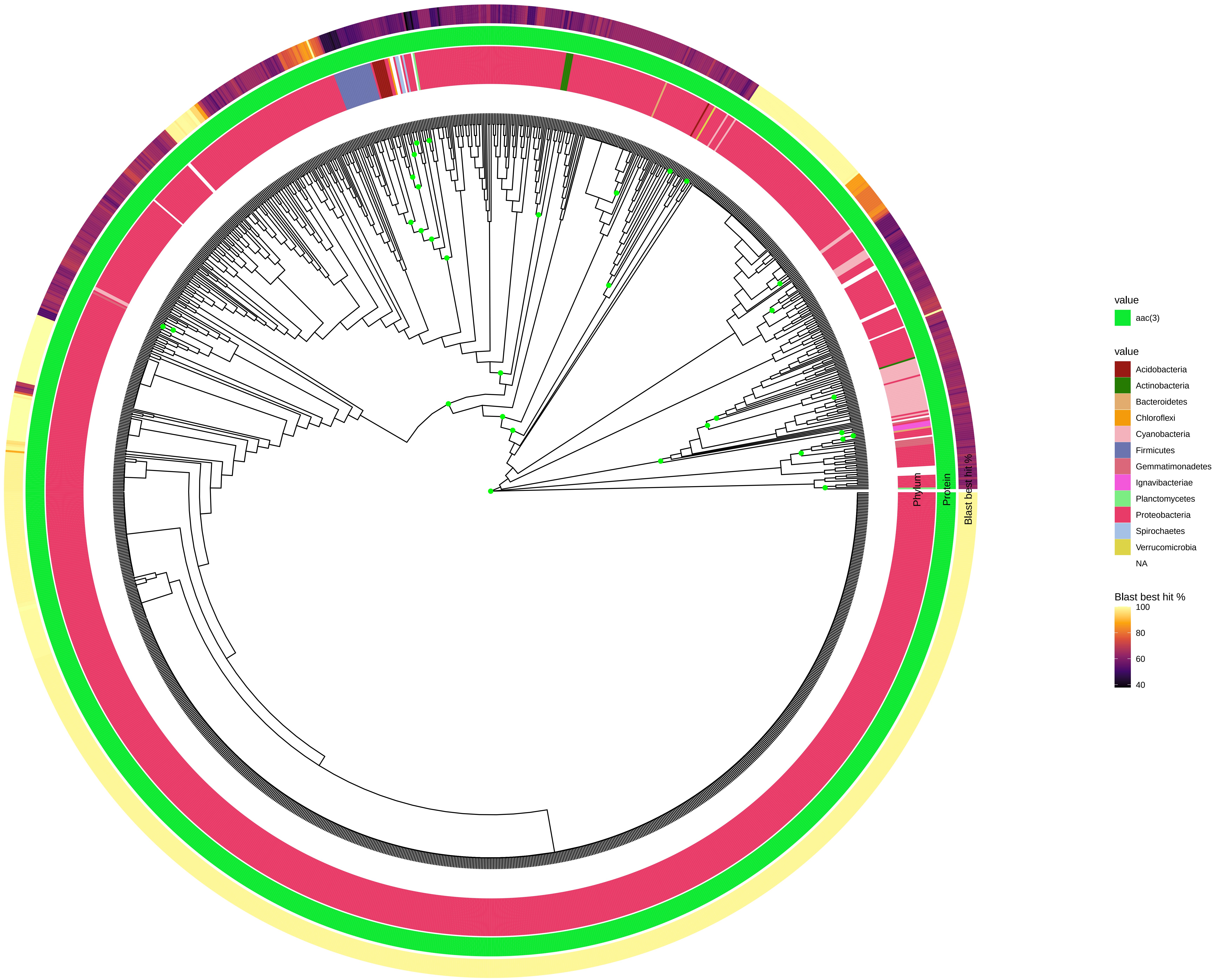
Phylum
Protein
Blast best hit %
value
aac(3)
value
Acidobacteria
Actinobacteria
Bacteroidetes
Chloroflexi
Cyanobacteria
Firmicutes
Gemmatimonadetes
Ignavibacteriae
Planctomycetes
Proteobacteria
Spirochaetes
Verrucomicrobia
NA
Blast best hit %
100
80
60
40

### aac3_class2

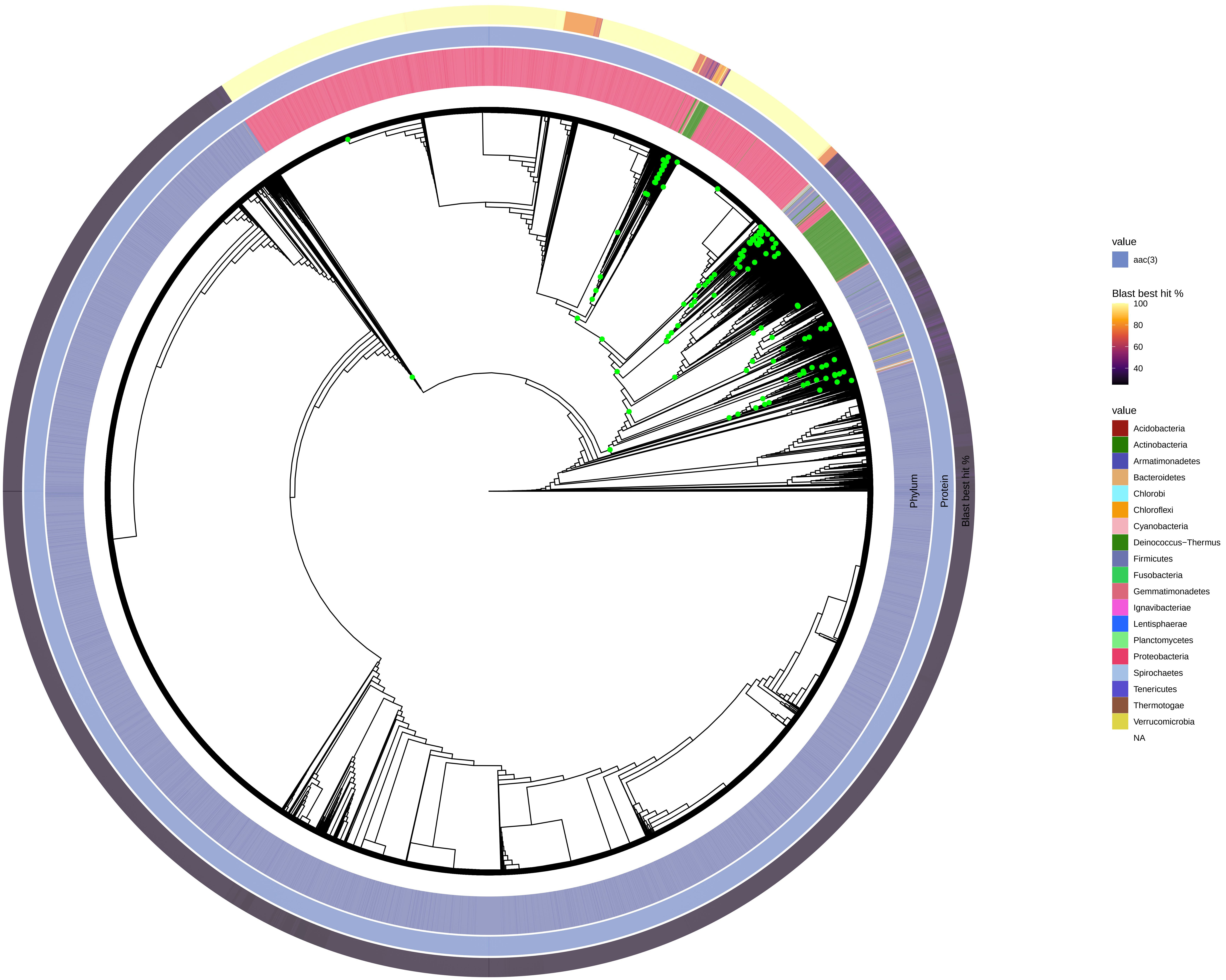

Phylum
Protein
Blast best hit %
value
aac(3)
Blast best hit %
100
80
60
40
value
Acidobacteria
Actinobacteria
Armatimonadetes
Bacteroidetes
Chlorobi
Chloroflexi
Cyanobacteria
Deinococcus−Thermus
Firmicutes
Fusobacteria
Gemmatimonadetes
Ignavibacteriae
Lentisphaerae
Planctomycetes
Proteobacteria
Spirochaetes
Tenericutes
Thermotogae
Verrucomicrobia
NA

### aac6p_complete

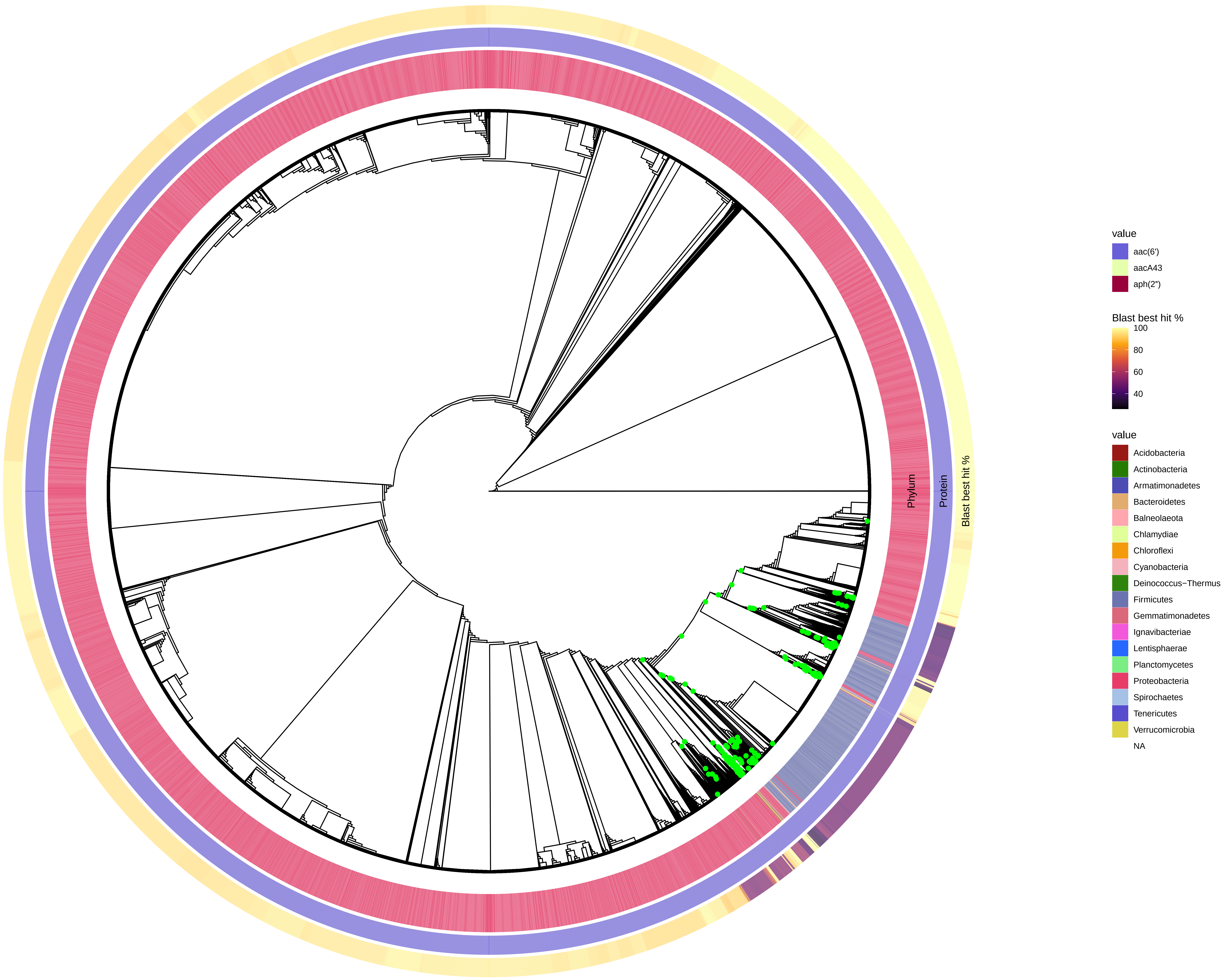

### aph2b

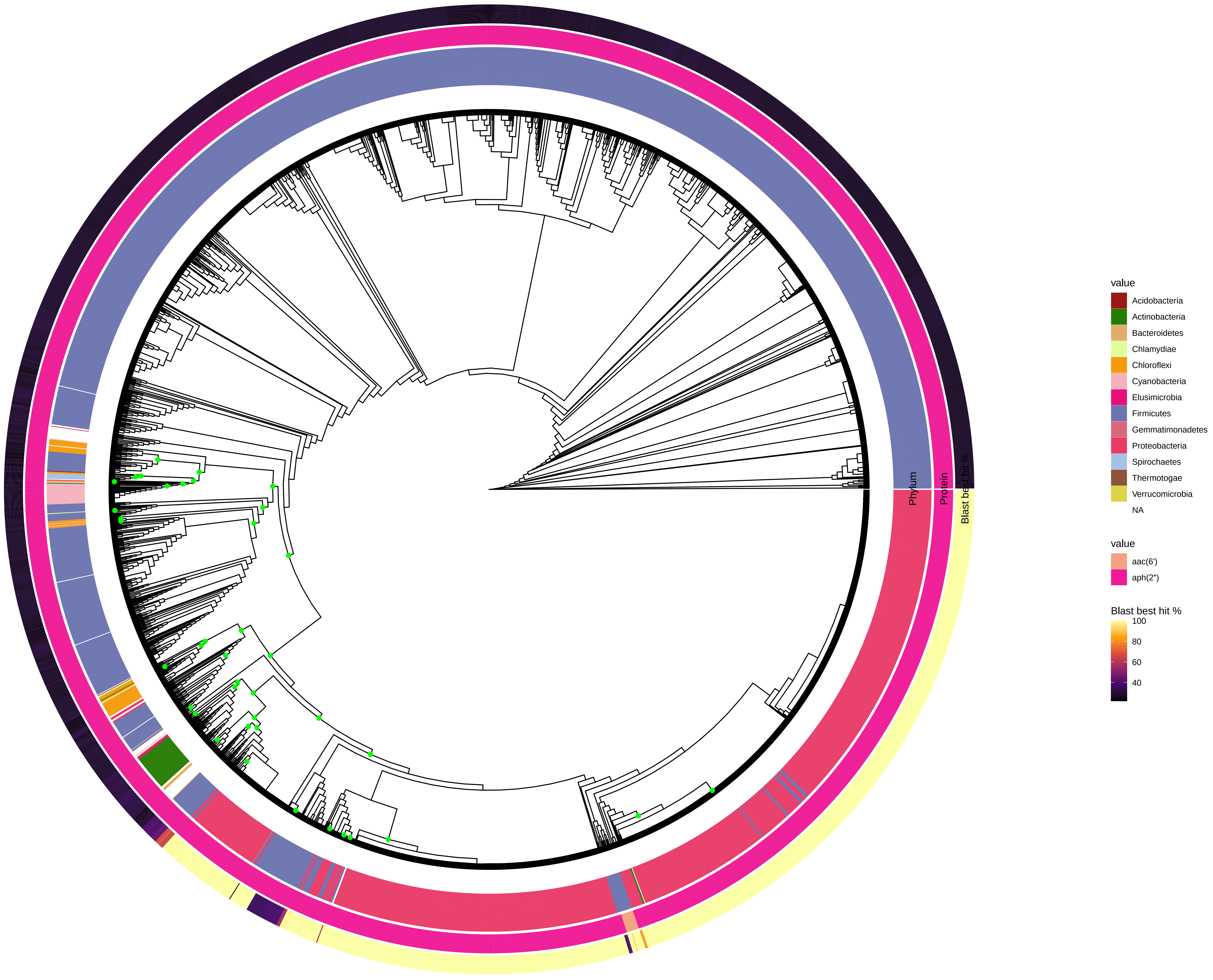

Phylum
Protein
Blast best hit %
value
Acidobacteria
Actinobacteria
Bacteroidetes
Chlamydiae
Chloroflexi
Cyanobacteria
Elusimicrobia
Firmicutes
Gemmatimonadetes
Proteobacteria
Spirochaetes
Thermotogae
Verrucomicrobia
NA
value
aac(6')
aph(2")
Blast best hit %
100
80
60
40

### aph3p

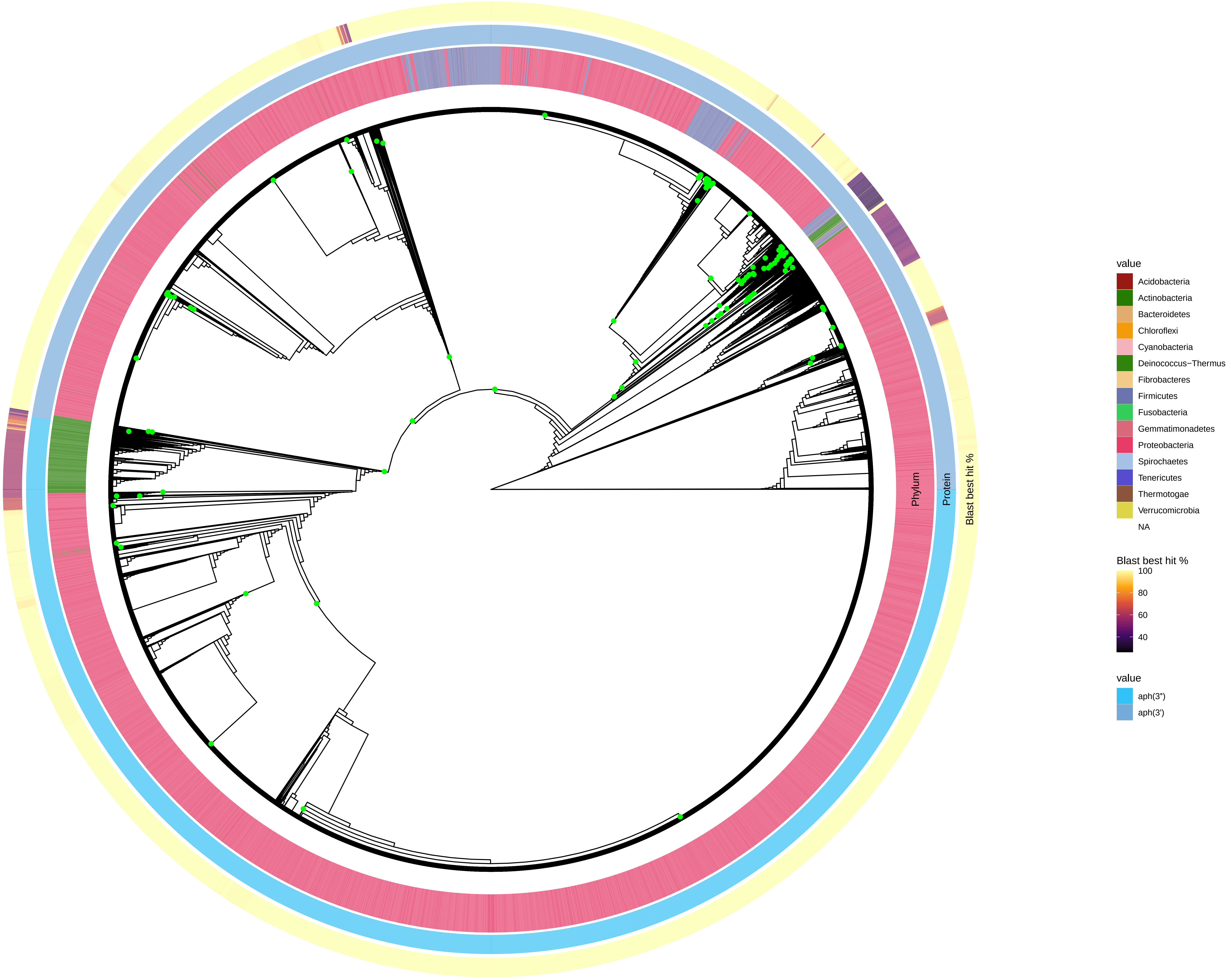

Phylum
Protein
Blast best hit %
value
Acidobacteria
Actinobacteria
Bacteroidetes
Chloroflexi
Cyanobacteria
Deinococcus−Thermus
Fibrobacteres
Firmicutes
Fusobacteria
Gemmatimonadetes
Proteobacteria
Spirochaetes
Tenericutes
Thermotogae
Verrucomicrobia
NA
Blast best hit %
100
80
60
40
value
aph(3")
aph(3')

### aph6

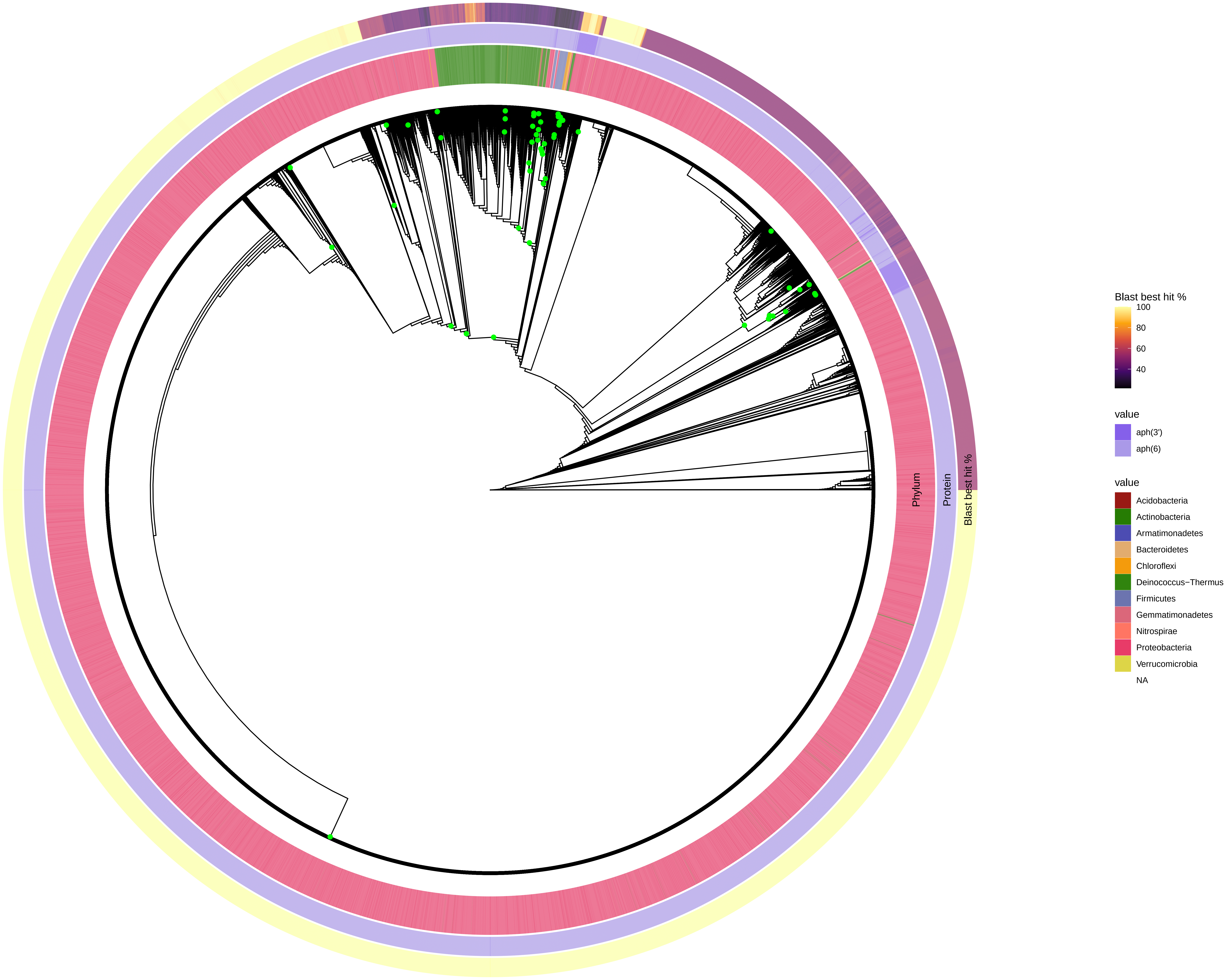

Phylum
Protein
Blast best hit %
Blast best hit %
100
80
60
40
value
aph(3')
aph(6)
value
Acidobacteria
Actinobacteria
Armatimonadetes
Bacteroidetes
Chloroflexi
Deinococcus−Thermus
Firmicutes
Gemmatimonadetes
Nitrospirae
Proteobacteria
Verrucomicrobia
NA

### class_a

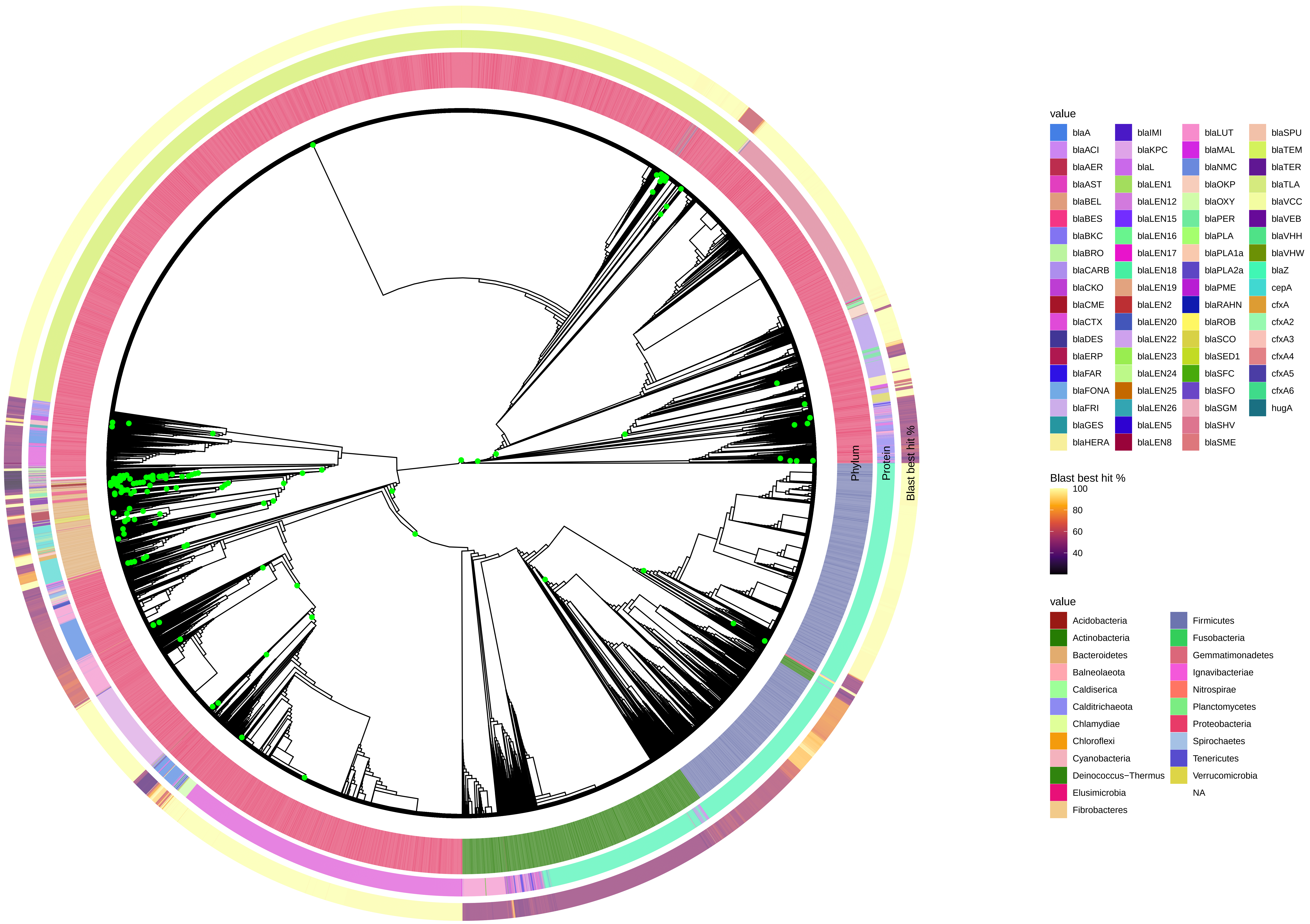

Phylum
Protein
Blast best hit %
value
blaA
blaACI
blaAER
blaAST
blaBEL
blaBES
blaBKC
blaBRO
blaCARB
blaCKO
blaCME
blaCTX
blaDES
blaERP
blaFAR
blaFONA
blaFRI
blaGES
blaHERA
blaIMI
blaKPC
blaL
blaLEN1
blaLEN12
blaLEN15
blaLEN16
blaLEN17
blaLEN18
blaLEN19
blaLEN2
blaLEN20
blaLEN22
blaLEN23
blaLEN24
blaLEN25
blaLEN26
blaLEN5
blaLEN8
blaLUT
blaMAL
blaNMC
blaOKP
blaOXY
blaPER
blaPLA
blaPLA1a
blaPLA2a
blaPME
blaRAHN
blaROB
blaSCO
blaSED1
blaSFC
blaSFO
blaSGM
blaSHV
blaSME
blaSPU
blaTEM
blaTER
blaTLA
blaVCC
blaVEB
blaVHH
blaVHW
blaZ
cepA
cfxA
cfxA2
cfxA3
cfxA4
cfxA5
cfxA6
hugA
Blast best hit %
100
80
60
40
value
Acidobacteria
Actinobacteria
Bacteroidetes
Balneolaeota
Caldiserica
Calditrichaeota
Chlamydiae
Chloroflexi
Cyanobacteria
Deinococcus−Thermus
Elusimicrobia
Fibrobacteres
Firmicutes
Fusobacteria
Gemmatimonadetes
Ignavibacteriae
Nitrospirae
Planctomycetes
Proteobacteria
Spirochaetes
Tenericutes
Verrucomicrobia
NA

### class_b_1_2

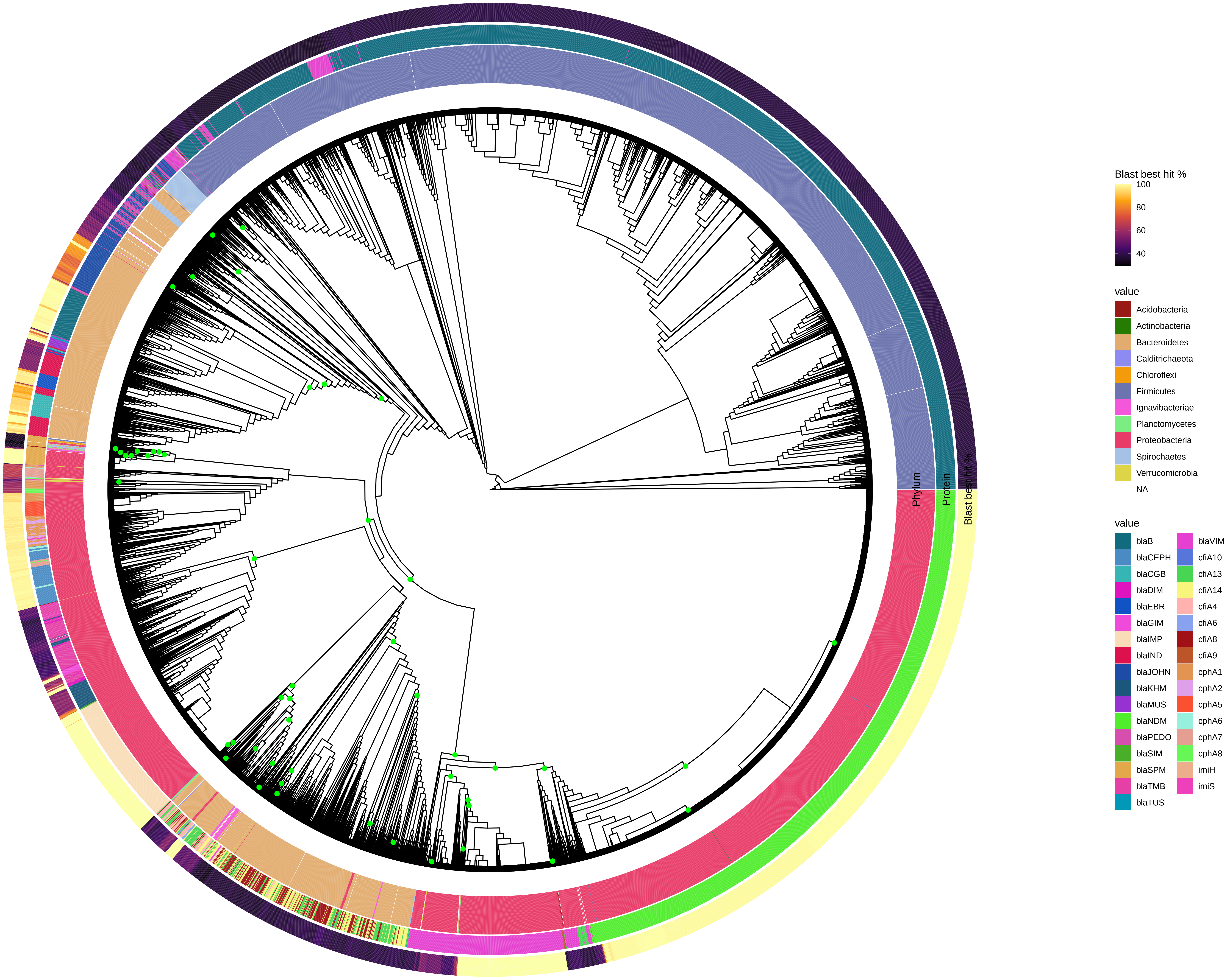

Phylum
Protein
Blast best hit %
Blast best hit %
100
80
60
40
value
Acidobacteria
Actinobacteria
Bacteroidetes
Calditrichaeota
Chloroflexi
Firmicutes
Ignavibacteriae
Planctomycetes
Proteobacteria
Spirochaetes
Verrucomicrobia
NA
value
blaB
blaCEPH
blaCGB
blaDIM
blaEBR
blaGIM
blaIMP
blaIND
blaJOHN
blaKHM
blaMUS
blaNDM
blaPEDO
blaSIM
blaSPM
blaTMB
blaTUS
blaVIM
cfiA10
cfiA13
cfiA14
cfiA4
cfiA6
cfiA8
cfiA9
cphA1
cphA2
cphA5
cphA6
cphA7
cphA8
imiH
imiS

### class_b_3

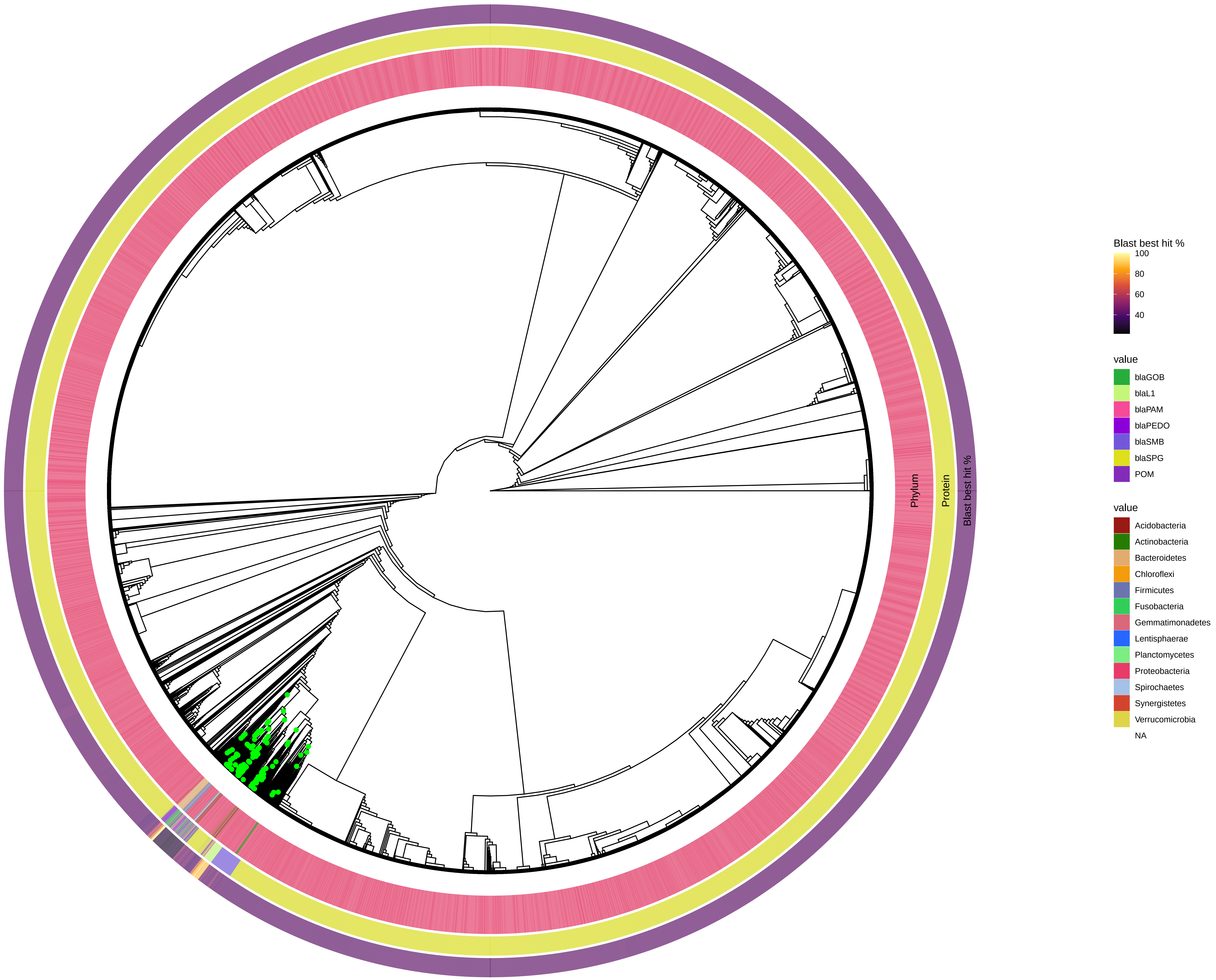

Phylum
Protein
Blast best hit %
Blast best hit %
100
80
60
40
value
blaGOB
blaL1
blaPAM
blaPEDO
blaSMB
blaSPG
POM
value
Acidobacteria
Actinobacteria
Bacteroidetes
Chloroflexi
Firmicutes
Fusobacteria
Gemmatimonadetes
Lentisphaerae
Planctomycetes
Proteobacteria
Spirochaetes
Synergistetes
Verrucomicrobia
NA

### class_c

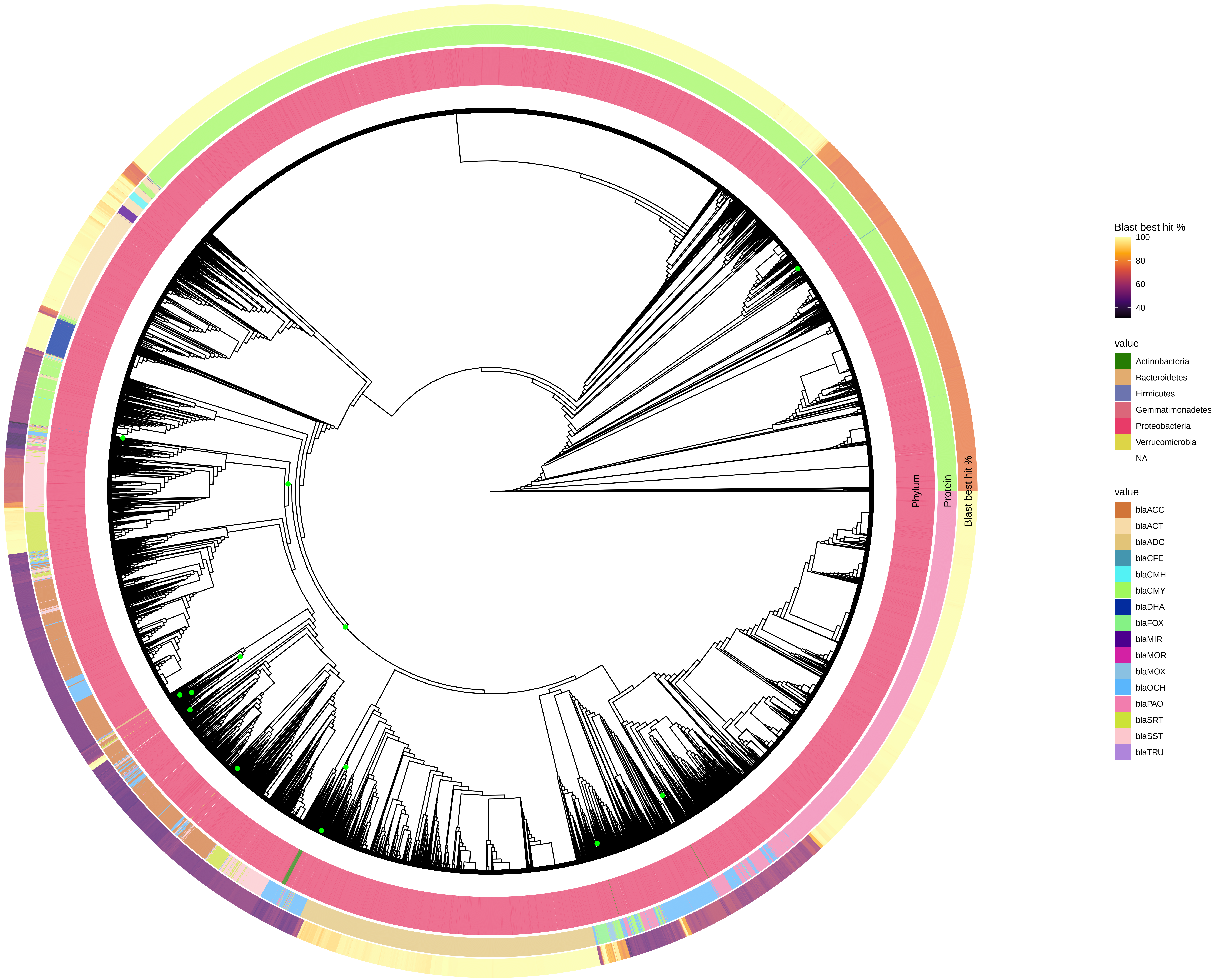
Phylum
Protein
Blast best hit %
Blast best hit %
100
80
60
40
value
Actinobacteria
Bacteroidetes
Firmicutes
Gemmatimonadetes
Proteobacteria
Verrucomicrobia
NA
value
blaACC
blaACT
blaADC
blaCFE
blaCMH
blaCMY
blaDHA
blaFOX
blaMIR
blaMOR
blaMOX
blaOCH
blaPAO
blaSRT
blaSST
blaTRU

### class_d_1_2

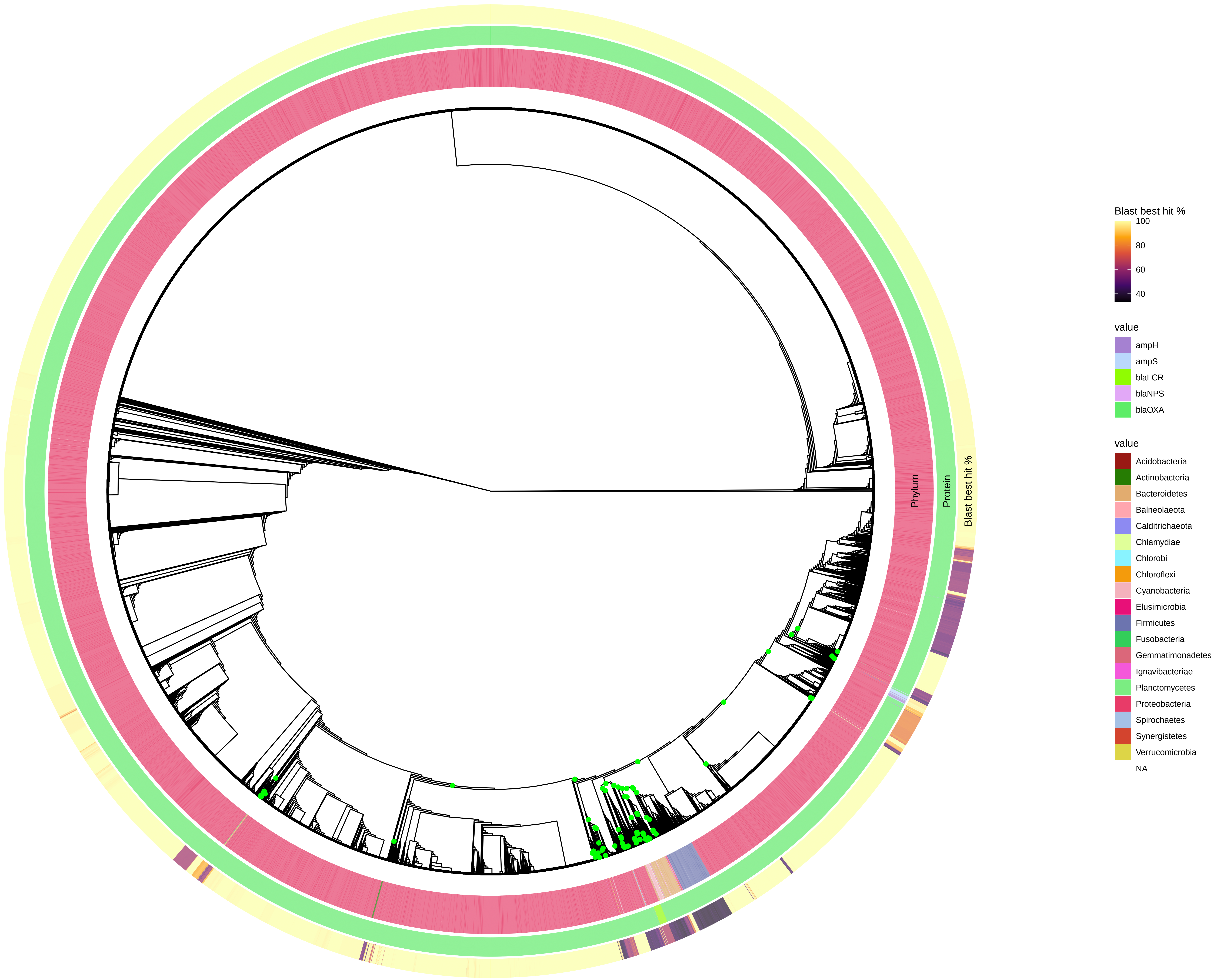
Phylum
Protein
Blast best hit %
Blast best hit %
100
80
60
40
value
ampH
ampS
blaLCR
blaNPS
blaOXA
value
Acidobacteria
Actinobacteria
Bacteroidetes
Balneolaeota
Calditrichaeota
Chlamydiae
Chlorobi
Chloroflexi
Cyanobacteria
Elusimicrobia
Firmicutes
Fusobacteria
Gemmatimonadetes
Ignavibacteriae
Planctomycetes
Proteobacteria
Spirochaetes
Synergistetes
Verrucomicrobia
NA

### macro_phospho

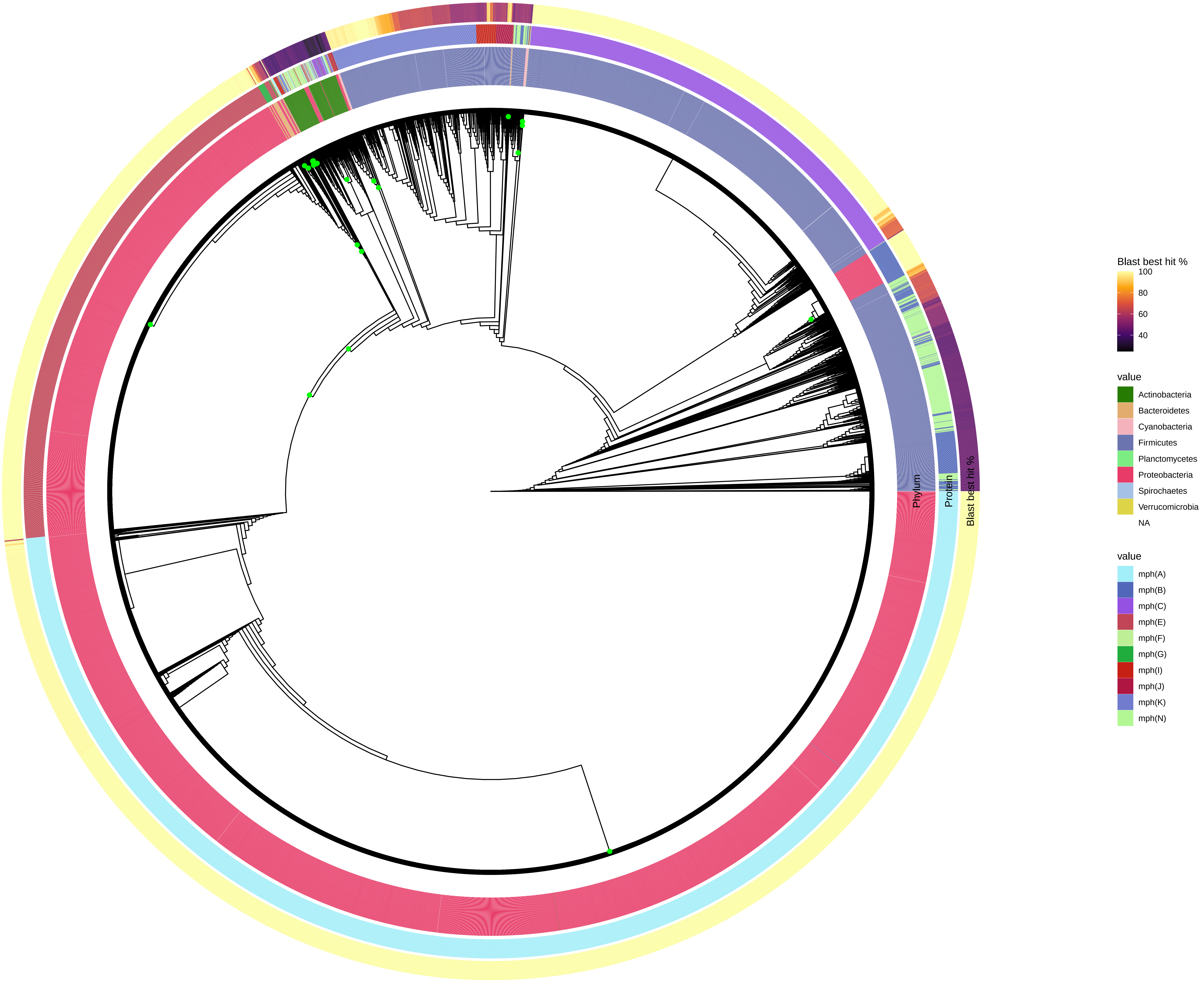

### methyltransf

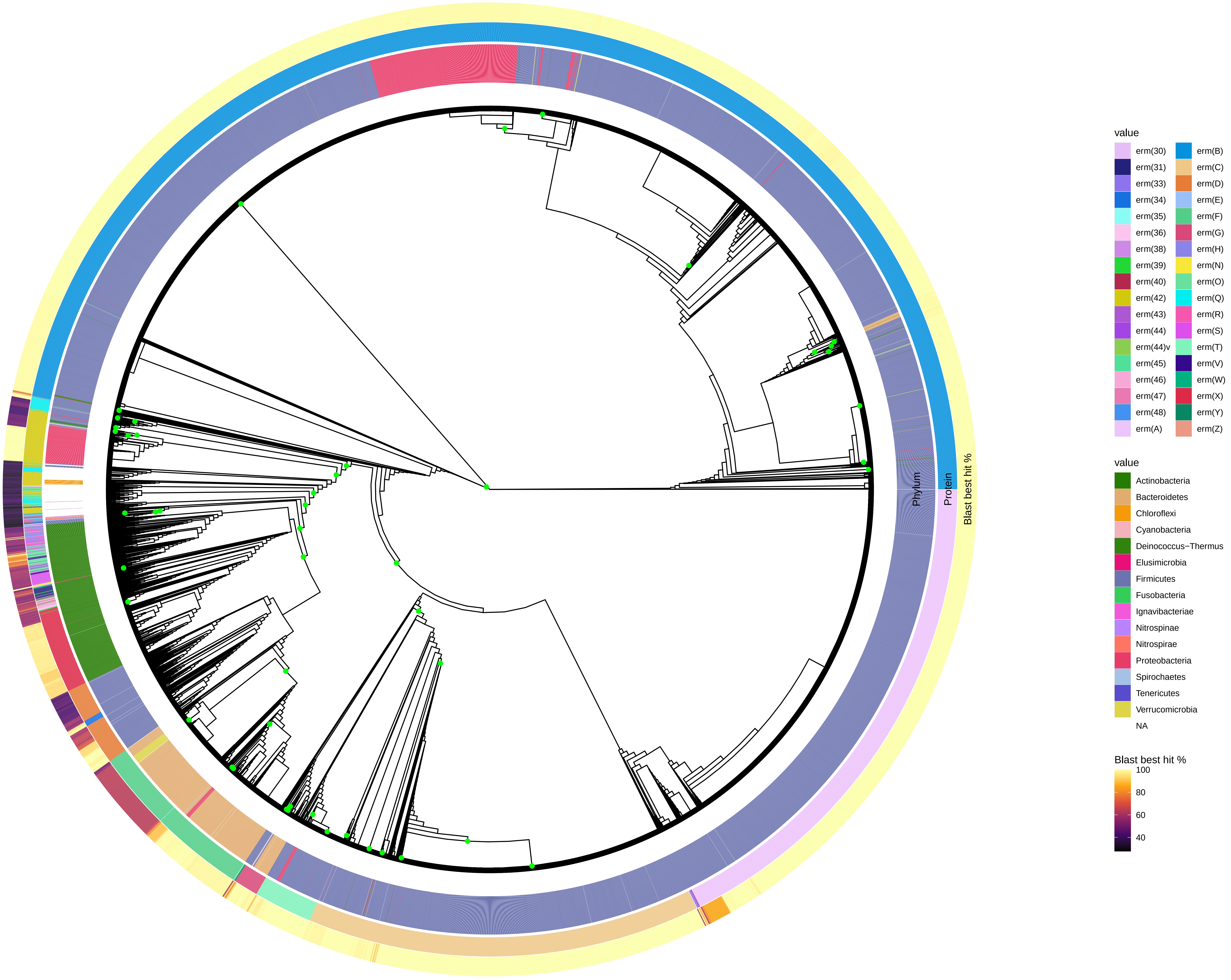

### qnr

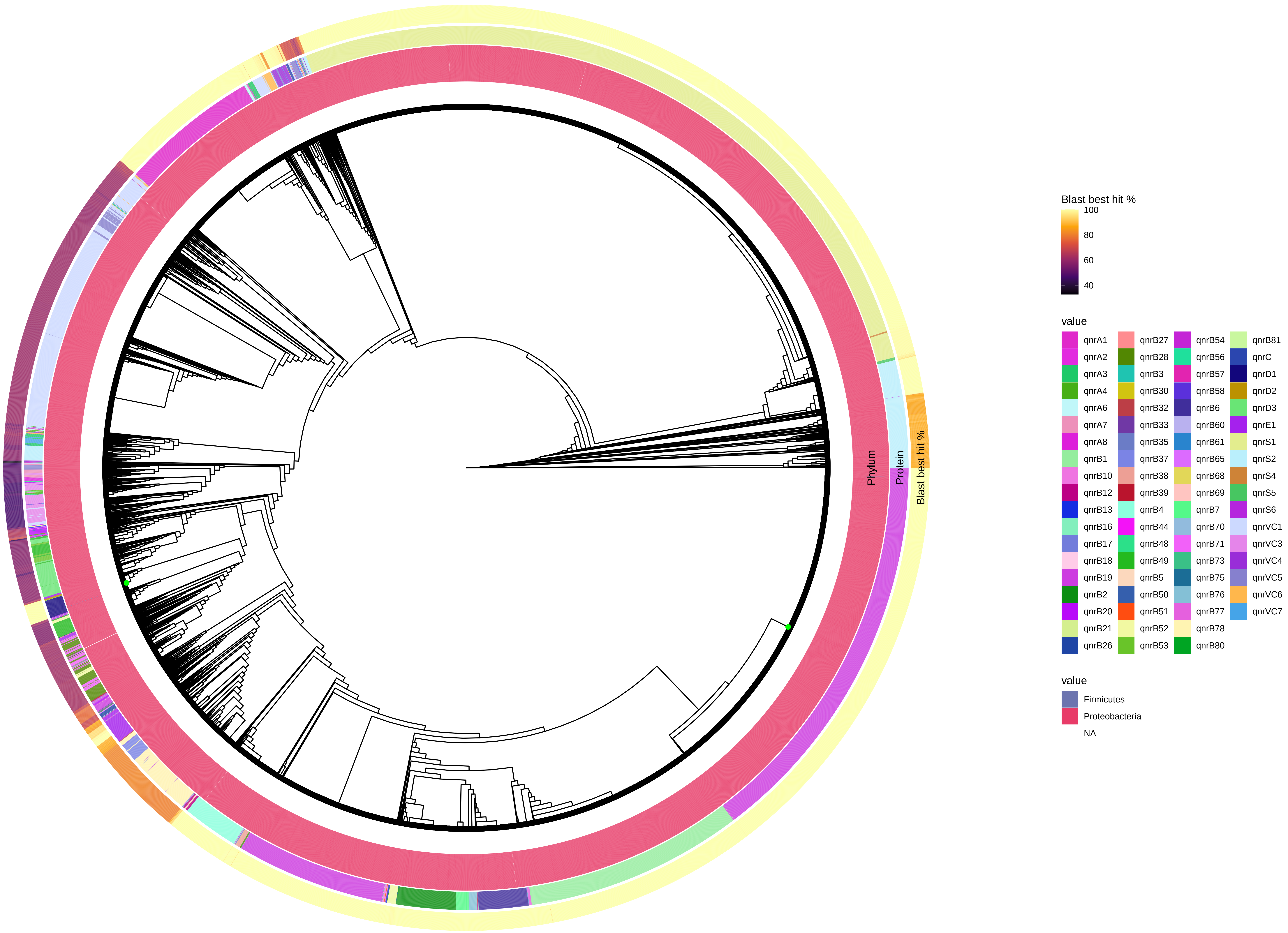

Phylum
Protein
Blast best hit %
Blast best hit %
100
80
60
40
value
qnrA1
qnrA2
qnrA3
qnrA4
qnrA6
qnrA7
qnrA8
qnrB1
qnrB10
qnrB12
qnrB13
qnrB16
qnrB17
qnrB18
qnrB19
qnrB2
qnrB20
qnrB21
qnrB26
qnrB27
qnrB28
qnrB3
qnrB30
qnrB32
qnrB33
qnrB35
qnrB37
qnrB38
qnrB39
qnrB4
qnrB44
qnrB48
qnrB49
qnrB5
qnrB50
qnrB51
qnrB52
qnrB53
qnrB54
qnrB56
qnrB57
qnrB58
qnrB6
qnrB60
qnrB61
qnrB65
qnrB68
qnrB69
qnrB7
qnrB70
qnrB71
qnrB73
qnrB75
qnrB76
qnrB77
qnrB78
qnrB80
qnrB81
qnrC
qnrD1
qnrD2
qnrD3
qnrE1
qnrS1
qnrS2
qnrS4
qnrS5
qnrS6
qnrVC1
qnrVC3
qnrVC4
qnrVC5
qnrVC6
qnrVC7
value
Firmicutes
Proteobacteria
NA

### tet_efflux

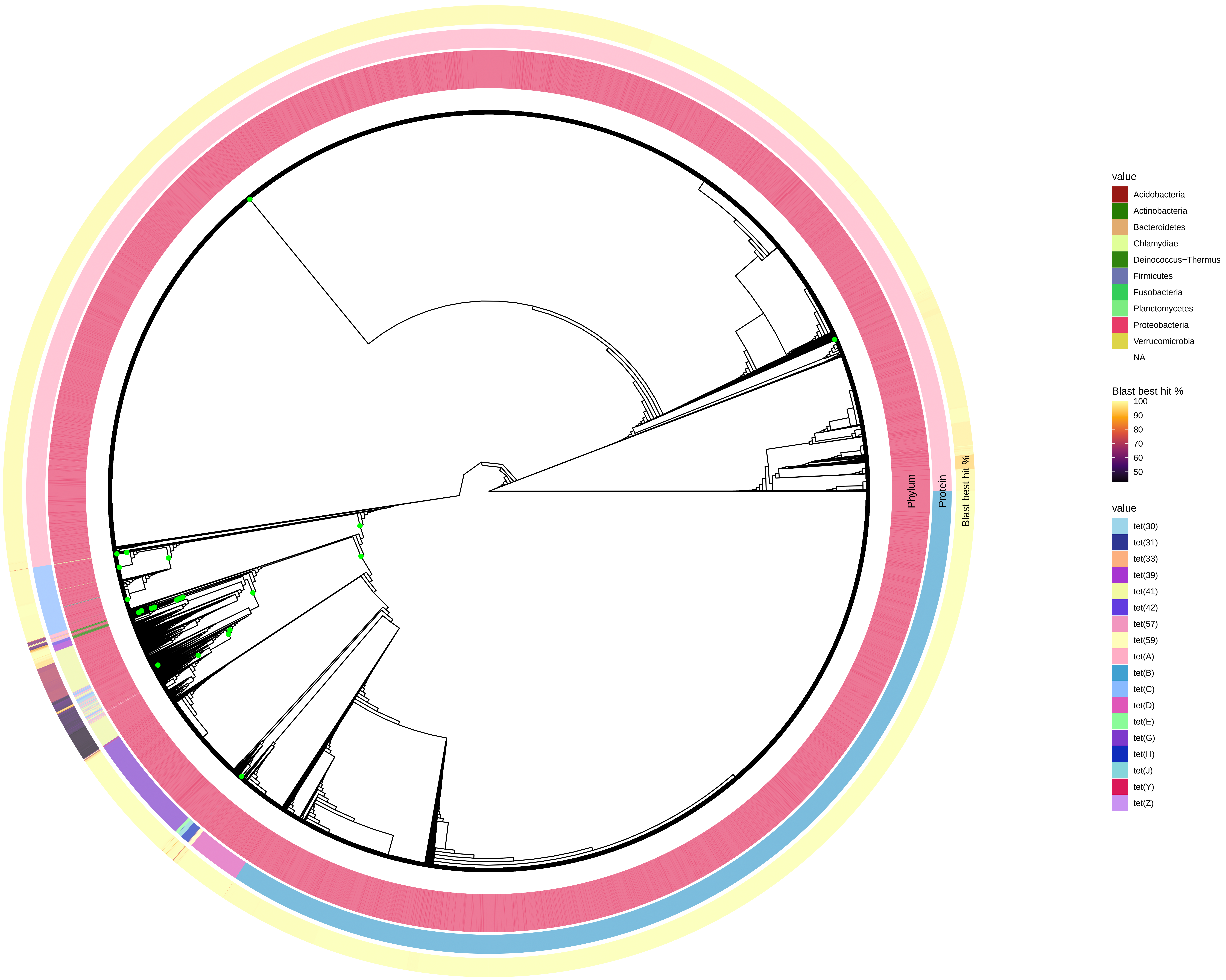

Phylum
Protein
Blast best hit %
value
Acidobacteria
Actinobacteria
Bacteroidetes
Chlamydiae
Deinococcus−Thermus
Firmicutes
Fusobacteria
Planctomycetes
Proteobacteria
Verrucomicrobia
NA
Blast best hit %
100
90
80
70
60
50
value
tet(30)
tet(31)
tet(33)
tet(39)
tet(41)
tet(42)
tet(57)
tet(59)
tet(A)
tet(B)
tet(C)
tet(D)
tet(E)
tet(G)
tet(H)
tet(J)
tet(Y)
tet(Z)

### tet_enzyme

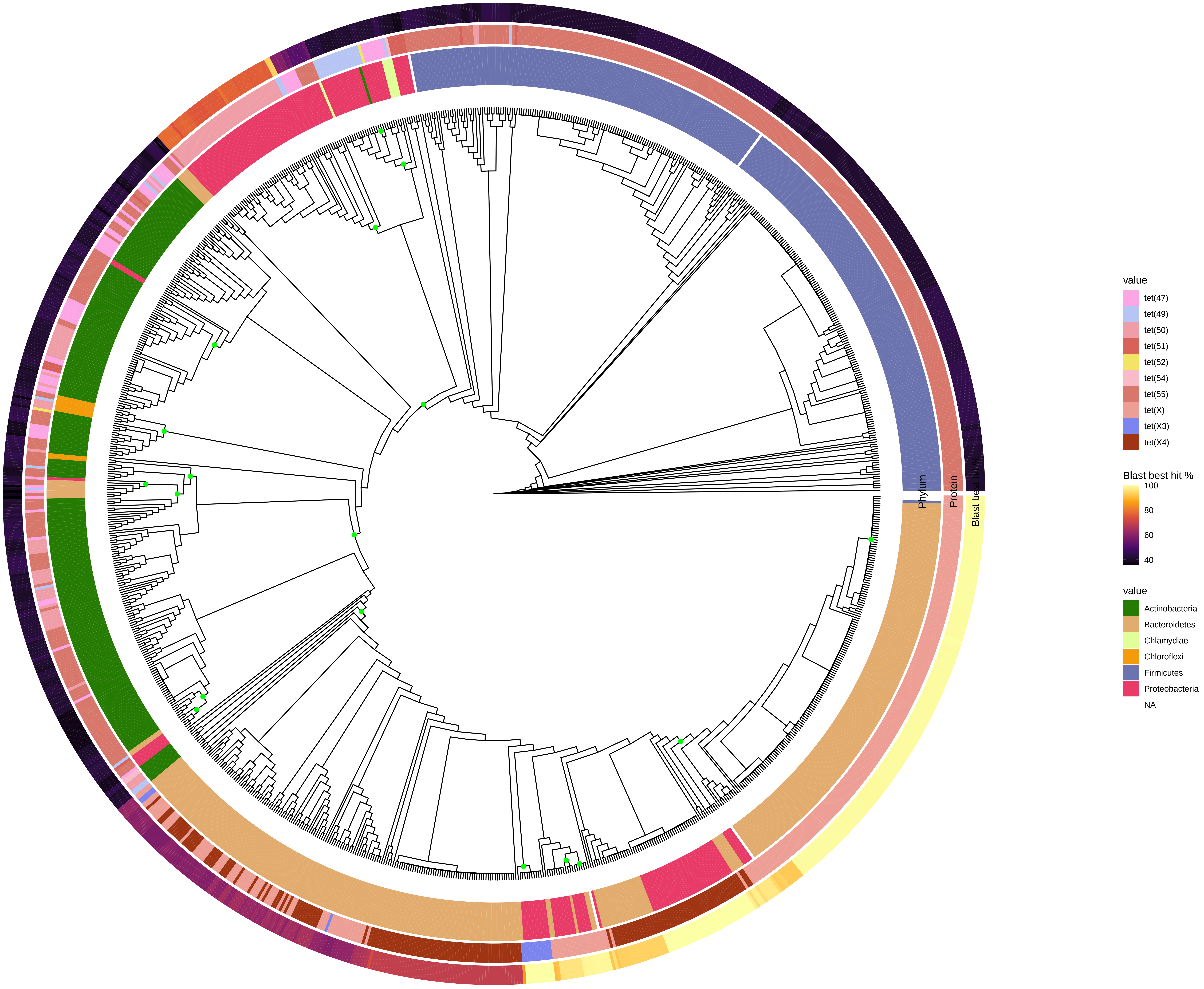

Phylum
Protein
Blast best hit %
value
tet(47)
tet(49)
tet(50)
tet(51)
tet(52)
tet(54)
tet(55)
tet(X)
tet(X3)
tet(X4)
Blast best hit %
100
80
60
40
value
Actinobacteria
Bacteroidetes
Chlamydiae
Chloroflexi
Firmicutes
Proteobacteria
NA

### tet_rpg

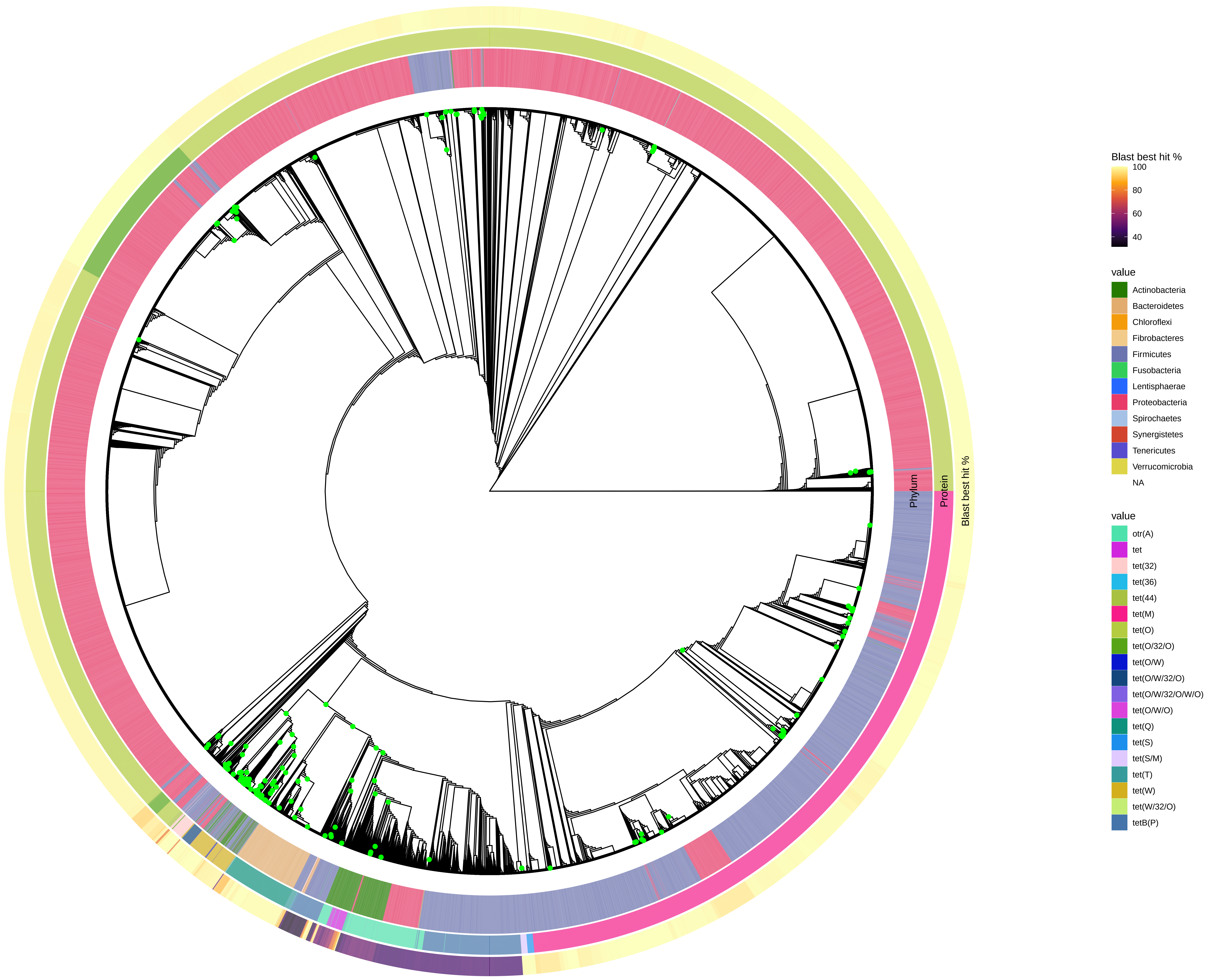

Phylum
Protein
Blast best hit %
Blast best hit %
100
80
60
40
value
Actinobacteria
Bacteroidetes
Chloroflexi
Fibrobacteres
Firmicutes
Fusobacteria
Lentisphaerae
Proteobacteria
Spirochaetes
Synergistetes
Tenericutes
Verrucomicrobia
NA
value
otr(A)
tet
tet(32)
tet(36)
tet(44)
tet(M)
tet(O)
tet(O/32/O)
tet(O/W)
tet(O/W/32/O)
tet(O/W/32/O/W/O)
tet(O/W/O)
tet(Q)
tet(S)
tet(S/M)
tet(T)
tet(W)
tet(W/32/O)
tetB(P)
