## Supplementary figures and images for "The transfer of antibiotic resistance genes between evolutionary distant bacteria"

### S19 Fig

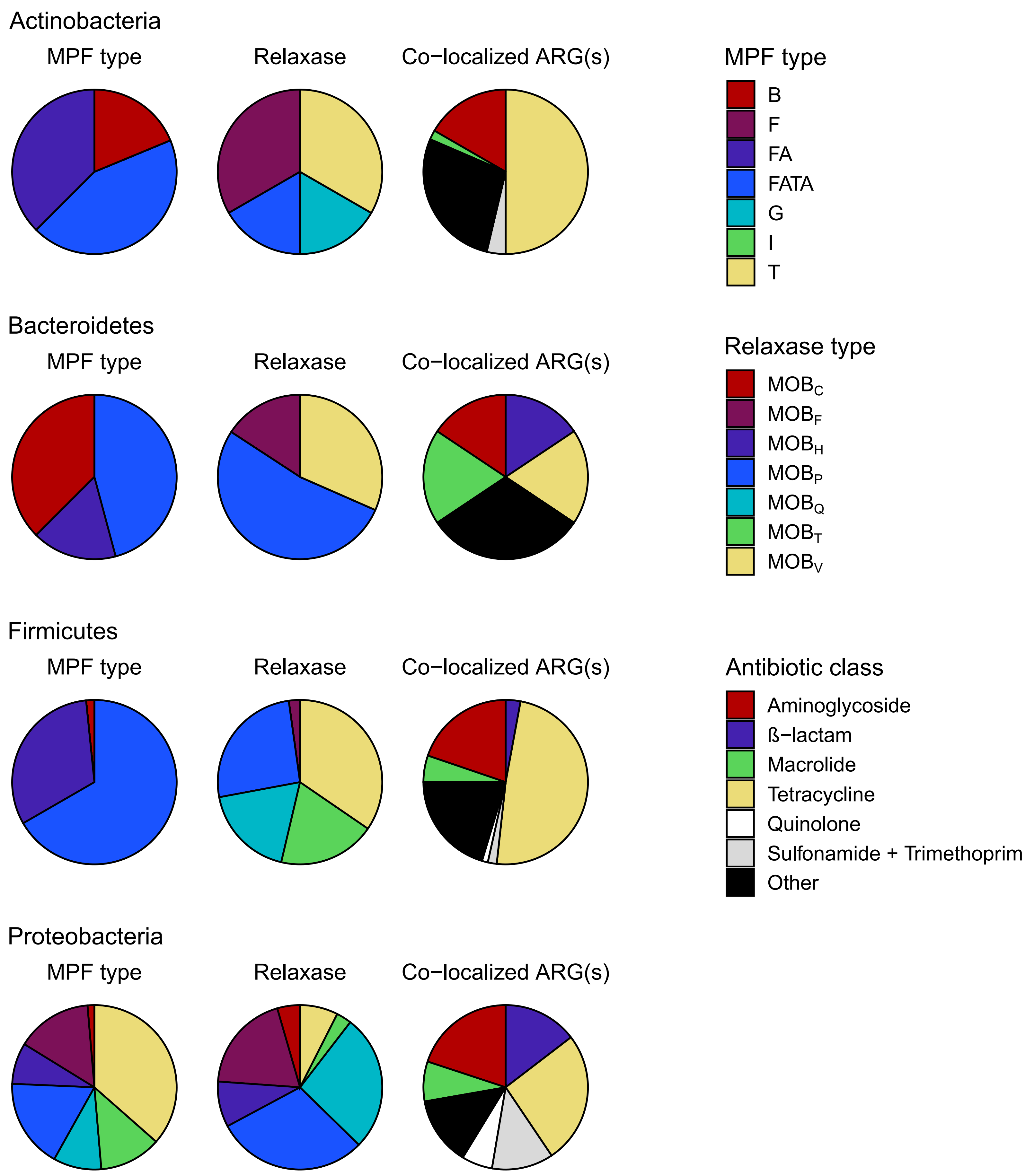
