## Supplementary material for "The transfer of antibiotic resistance genes between evolutionary distant bacteria": S20-37 Figs

aac2p

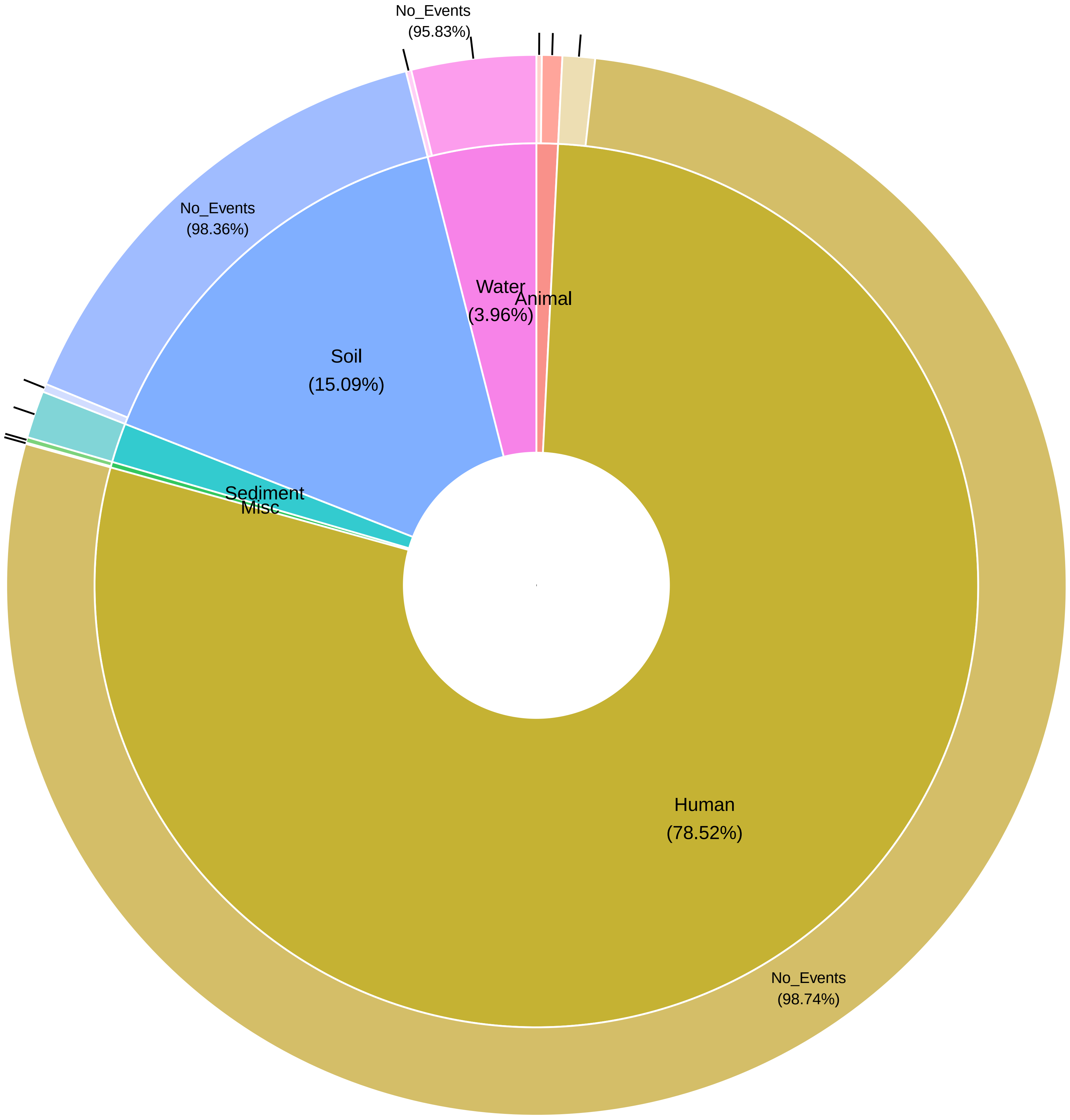
No_Events
(95.83%)
No_Events
(98.36%)
Water
(3.96%)
Animal
Soil
(15.09%)
Sediment
Misc
Human
(78.52%)
No_Events
(98.74%)

### aac3_class1

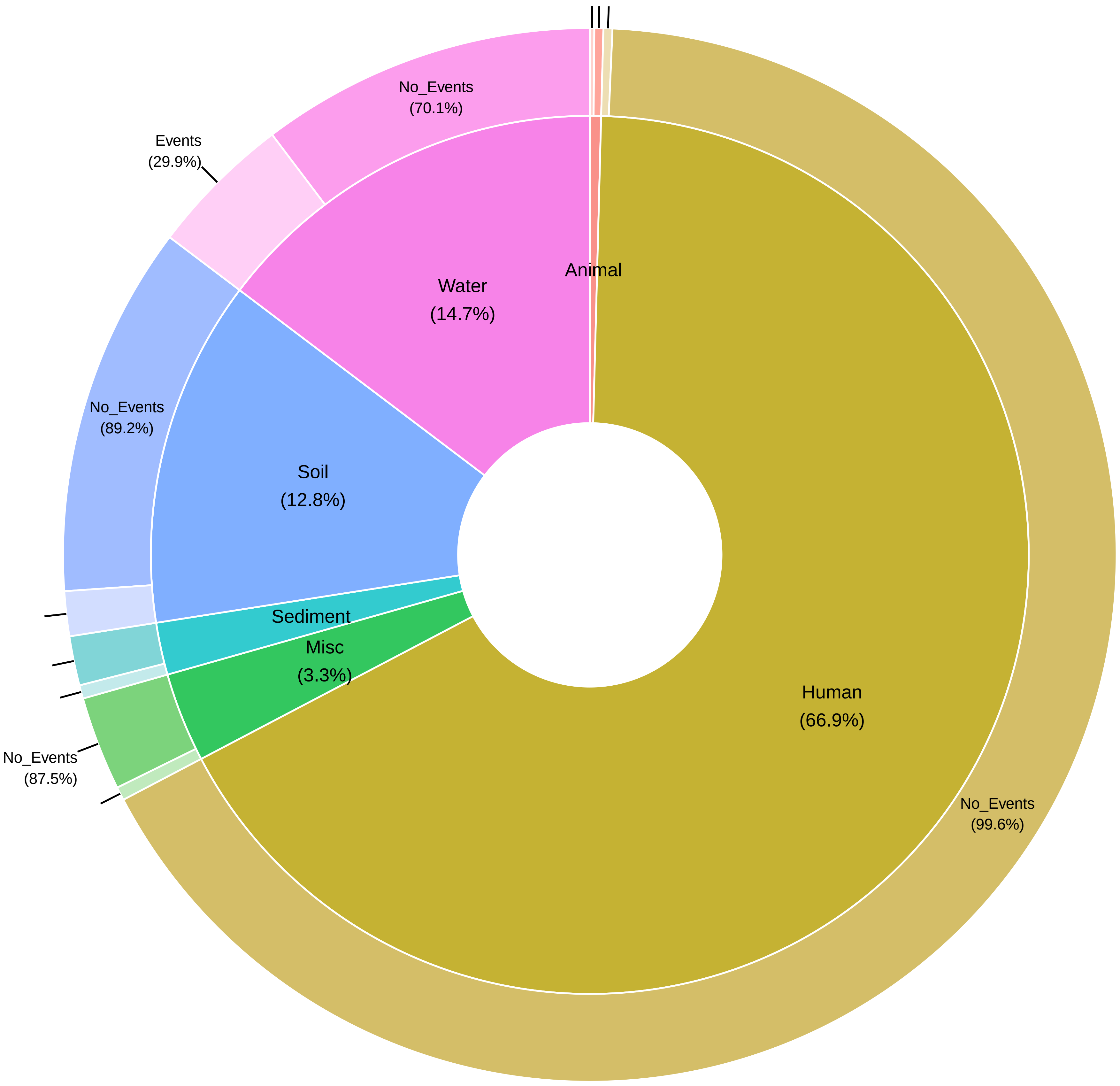

No_Events
(70.1%)
Events
(29.9%)
Water
(14.7%)
Animal
No_Events
(89.2%)
Soil
(12.8%)
Sediment
Misc
(3.3%)
Human
(66.9%)
No_Events
(87.5%)
No_Events
(99.6%)

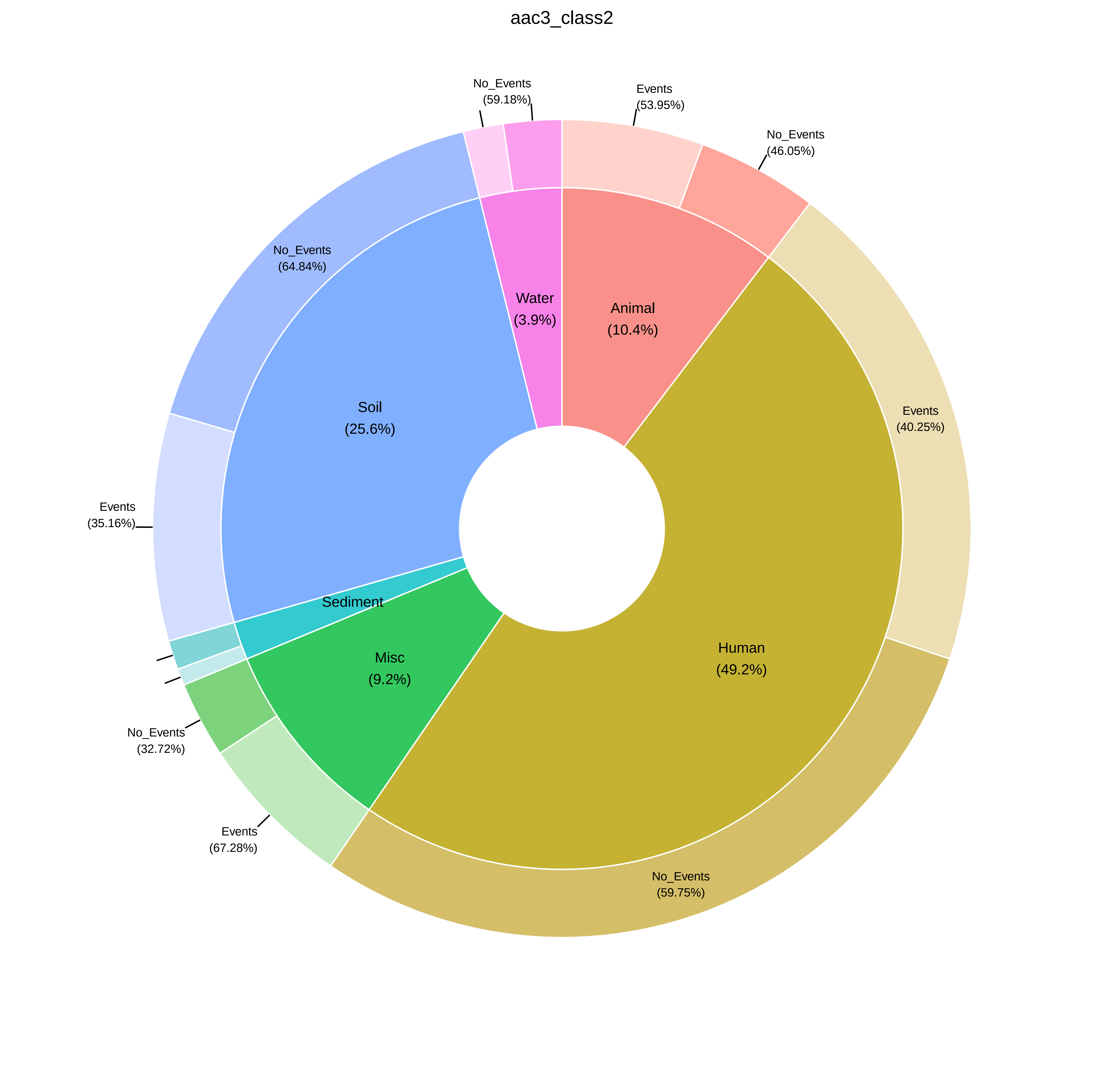

aac3_class2
No_Events
(59.18%)
Events
(53.95%)
No_Events
(46.05%)
No_Events
(64.84%)
Water
(3.9%)
Animal
(10.4%)
Soil
(25.6%)
Events
(40.25%)
Events
(35.16%)
Sediment
Misc
(9.2%)
Human
(49.2%)
No_Events
(32.72%)
Events
(67.28%)
No_Events
(59.75%)

aac6p_complete

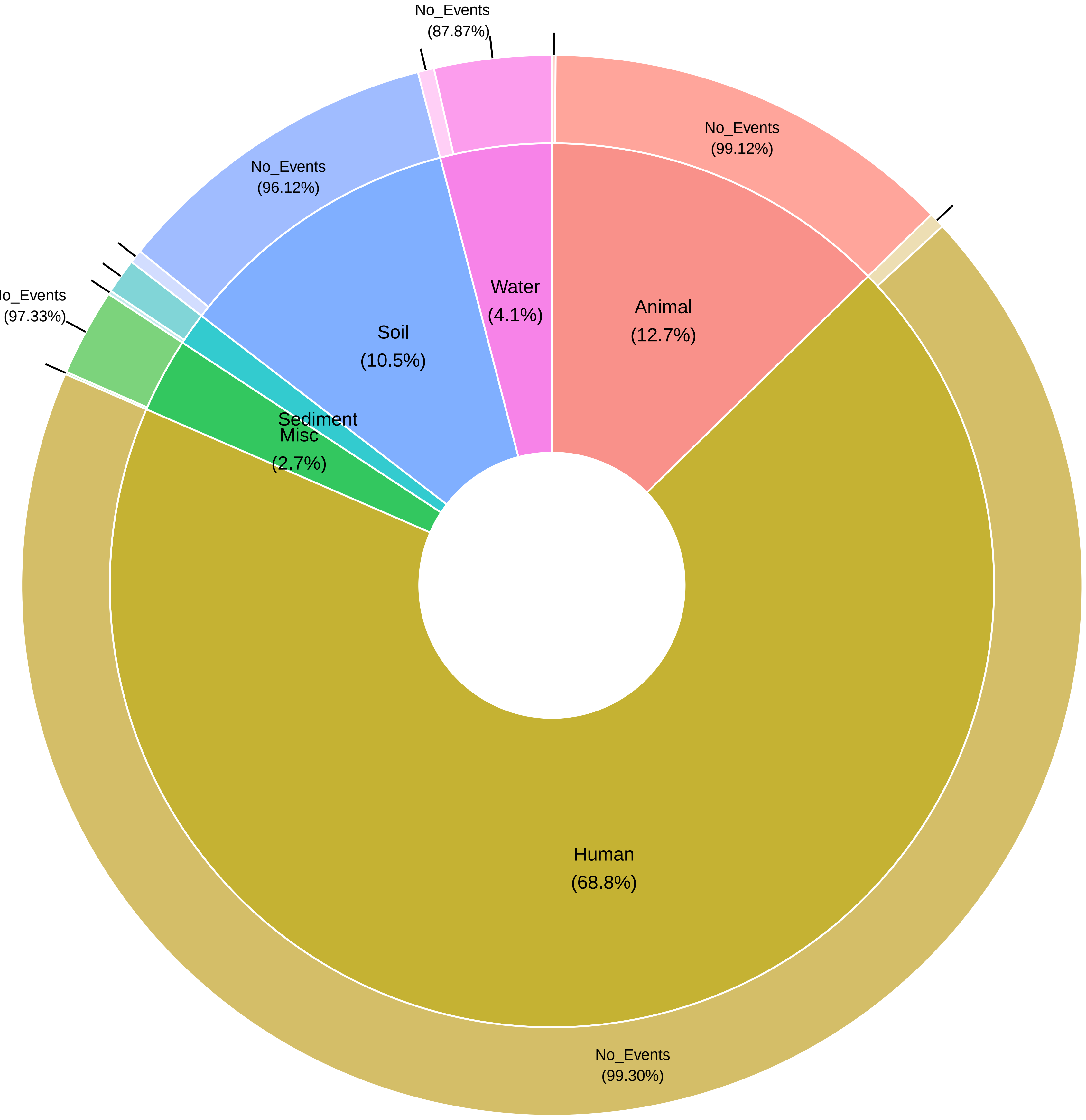

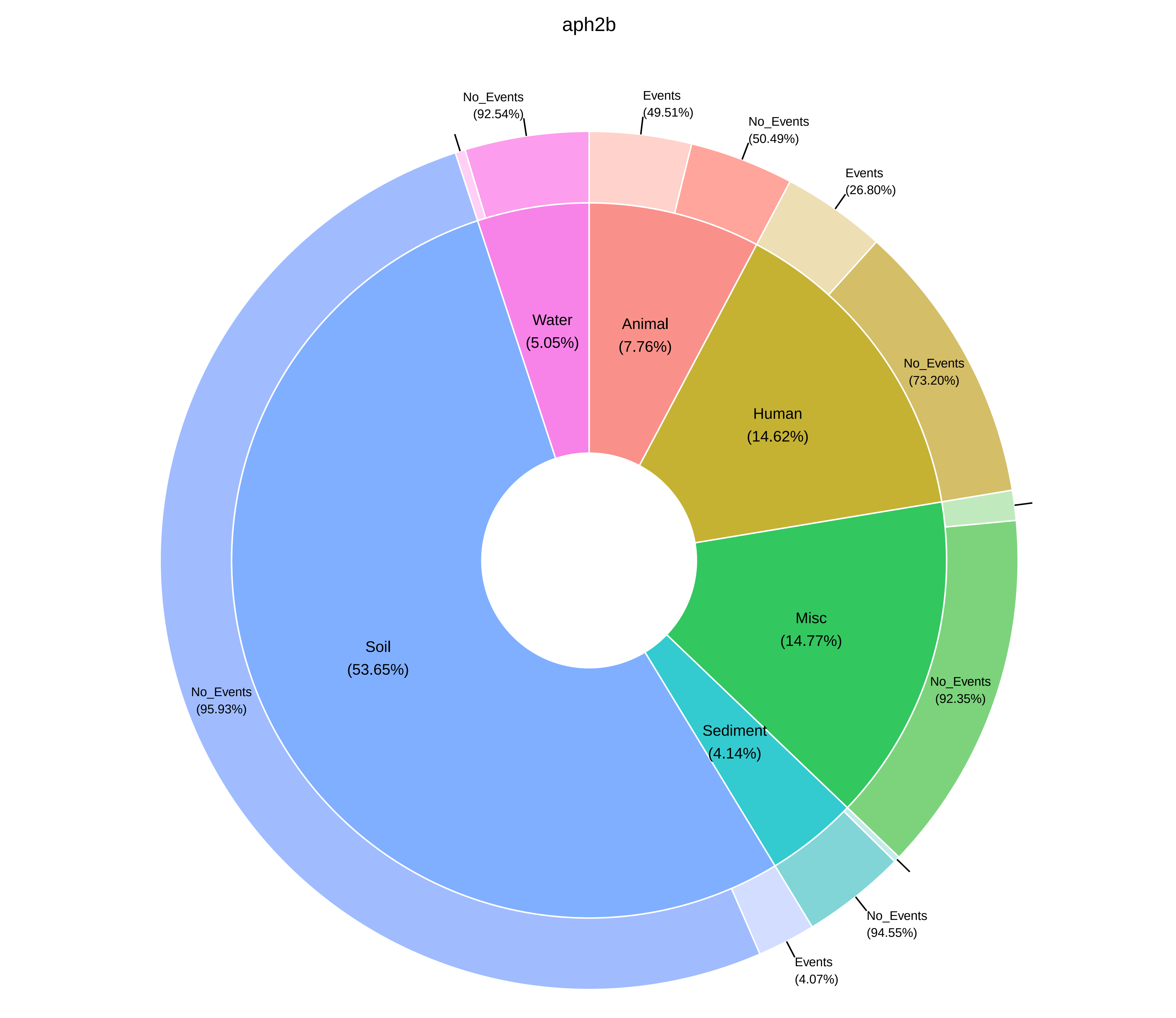
aph2b
No_Events
(92.54%)
Events
(49.51%)
No_Events
(50.49%)
Events
(26.80%)
Water
(5.05%)
Animal
(7.76%)
No_Events
(73.20%)
Human
(14.62%)
Misc
(14.77%)
Soil
(53.65%)
No_Events
(95.93%)
No_Events
(92.35%)
Sediment
(4.14%)
No_Events
(94.55%)
Events
(4.07%)

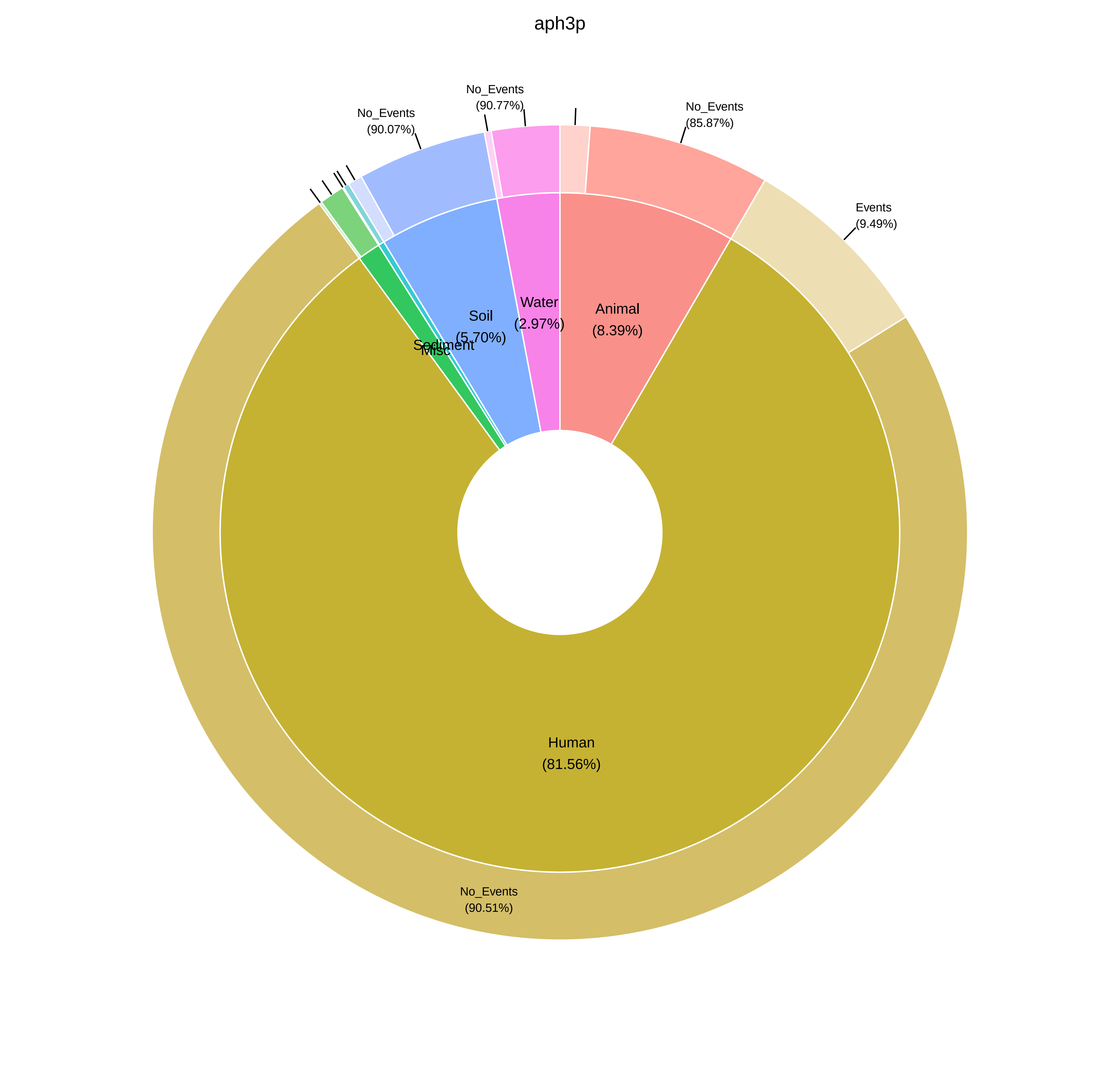

aph3p
No_Events
(90.07%)
No_Events
(90.77%)
No_Events
(85.87%)
Events
(9.49%)
Soil
(5.70%)
Water
(2.97%)
Animal
(8.39%)
Sediment
Misc
Human
(81.56%)
No_Events
(90.51%)

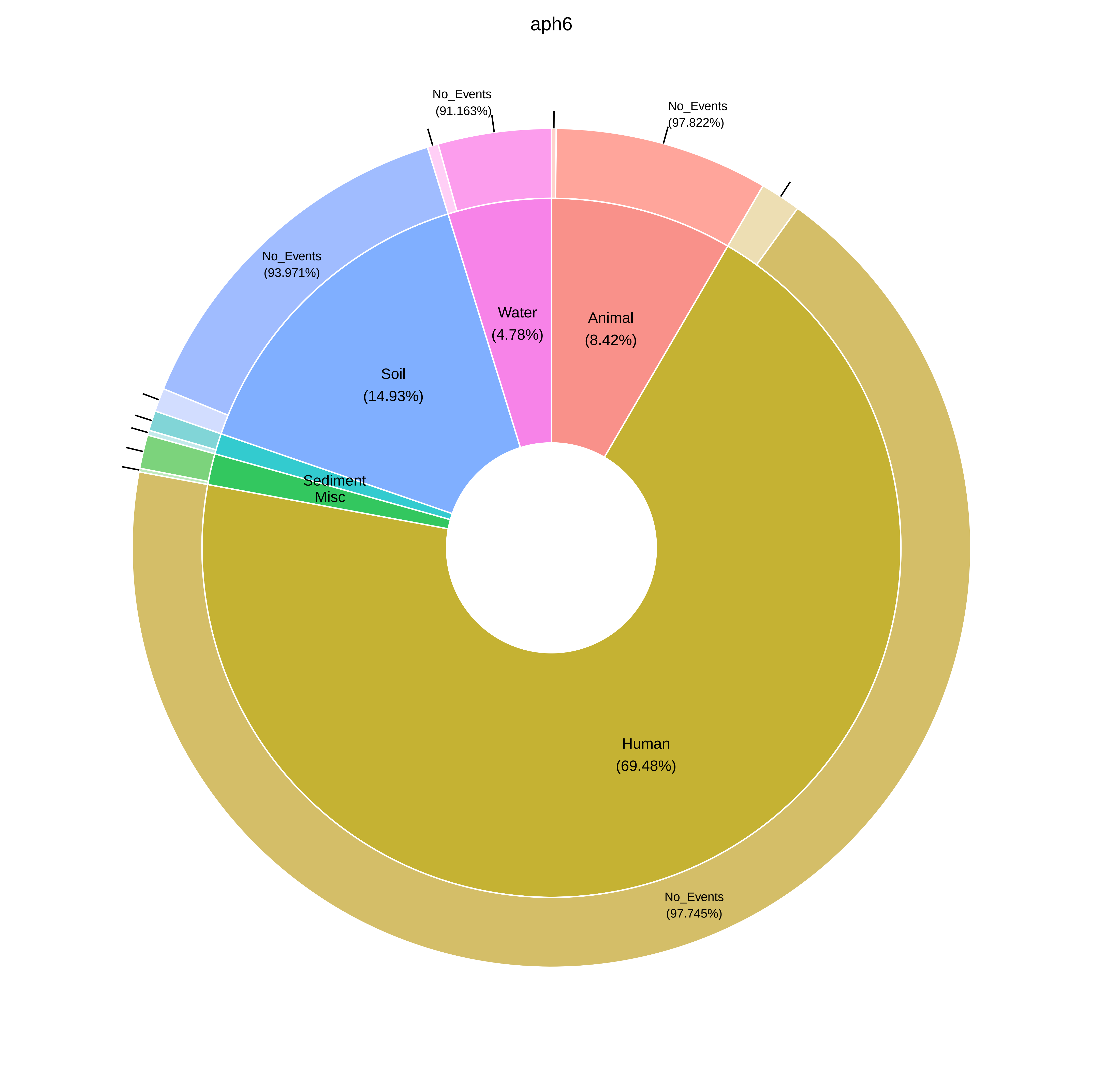
aph6
No_Events
(91.163%)
No_Events
(97.822%)
No_Events
(93.971%)
Water
(4.78%)
Animal
(8.42%)
Soil
(14.93%)
Sediment
Misc
Human
(69.48%)
No_Events
(97.745%)

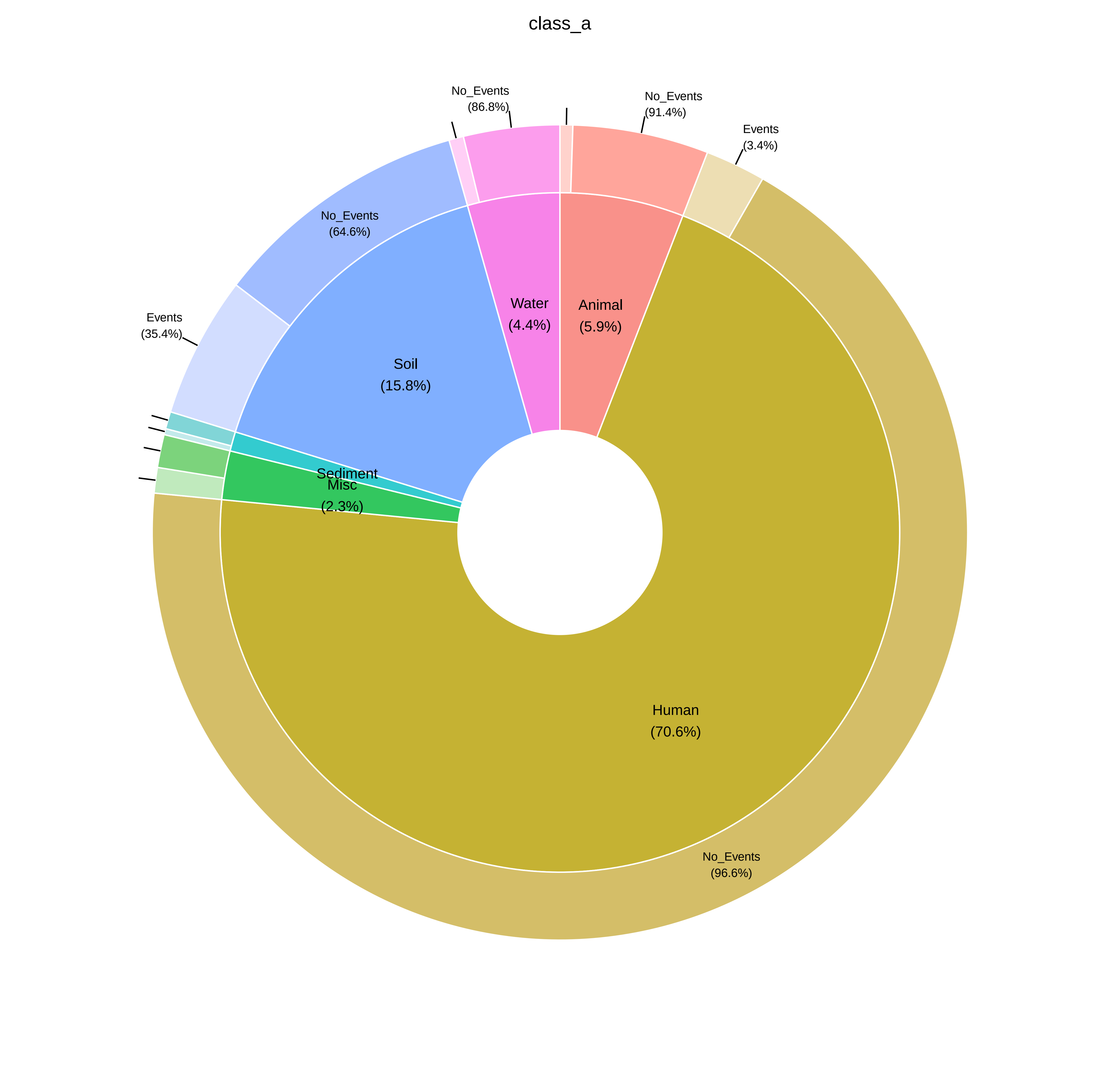
class_a
No_Events
(86.8%)
No_Events
(91.4%)
Events
(3.4%)
No_Events
(64.6%)
Events
(35.4%)
Water
(4.4%)
Animal
(5.9%)
Soil
(15.8%)
Sediment
Misc
(2.3%)
Human
(70.6%)
No_Events
(96.6%)

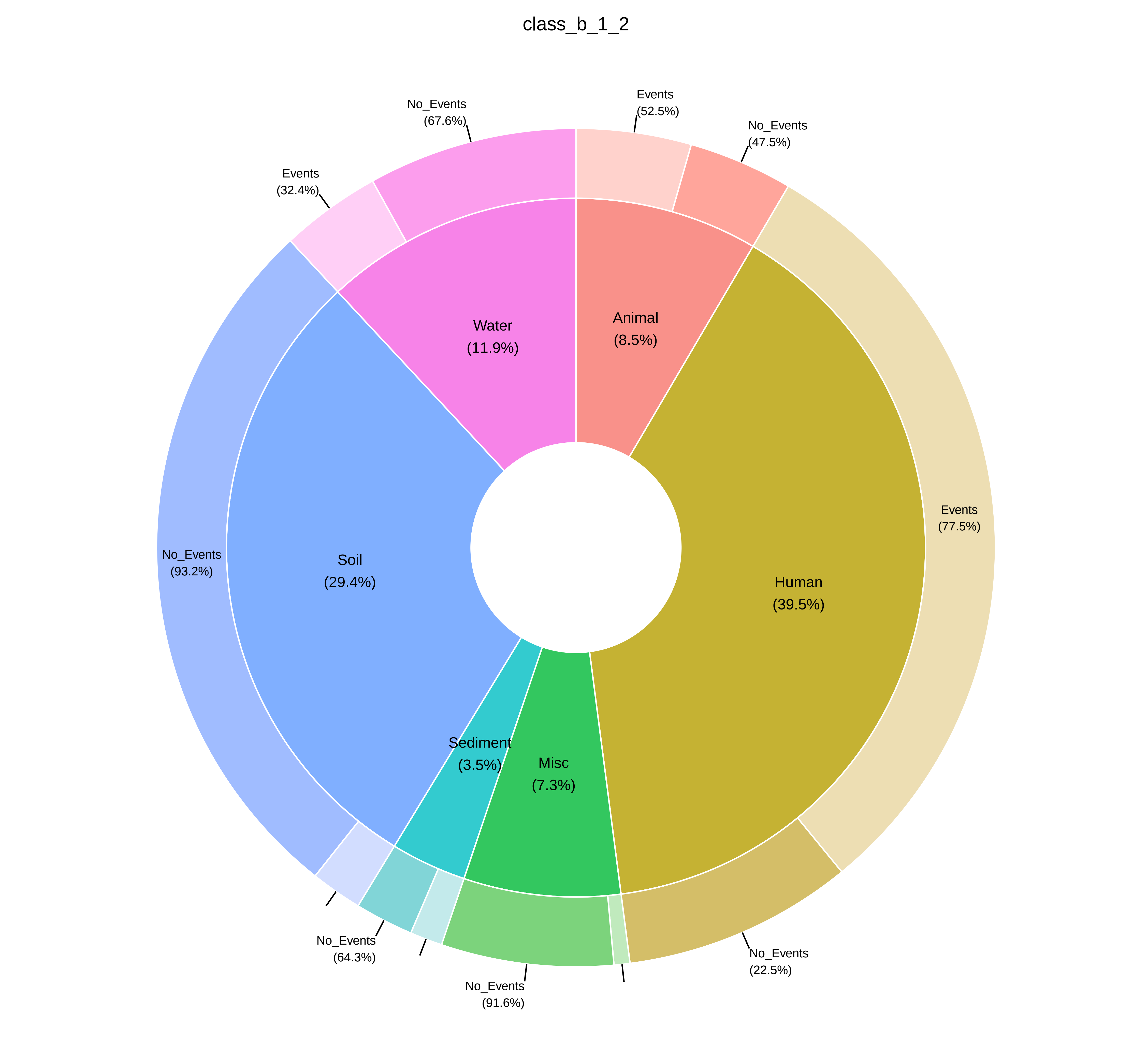
class_b_1_2
Water
(11.9%)
Animal
(8.5%)
Soil
(29.4%)
Human
(39.5%)
Sediment
(3.5%)
Misc
(7.3%)
No_Events
(67.6%)
Events
(52.5%)
No_Events
(47.5%)
Events
(32.4%)
Events
(77.5%)
No_Events
(93.2%)
No_Events
(64.3%)
No_Events
(91.6%)
No_Events
(22.5%)

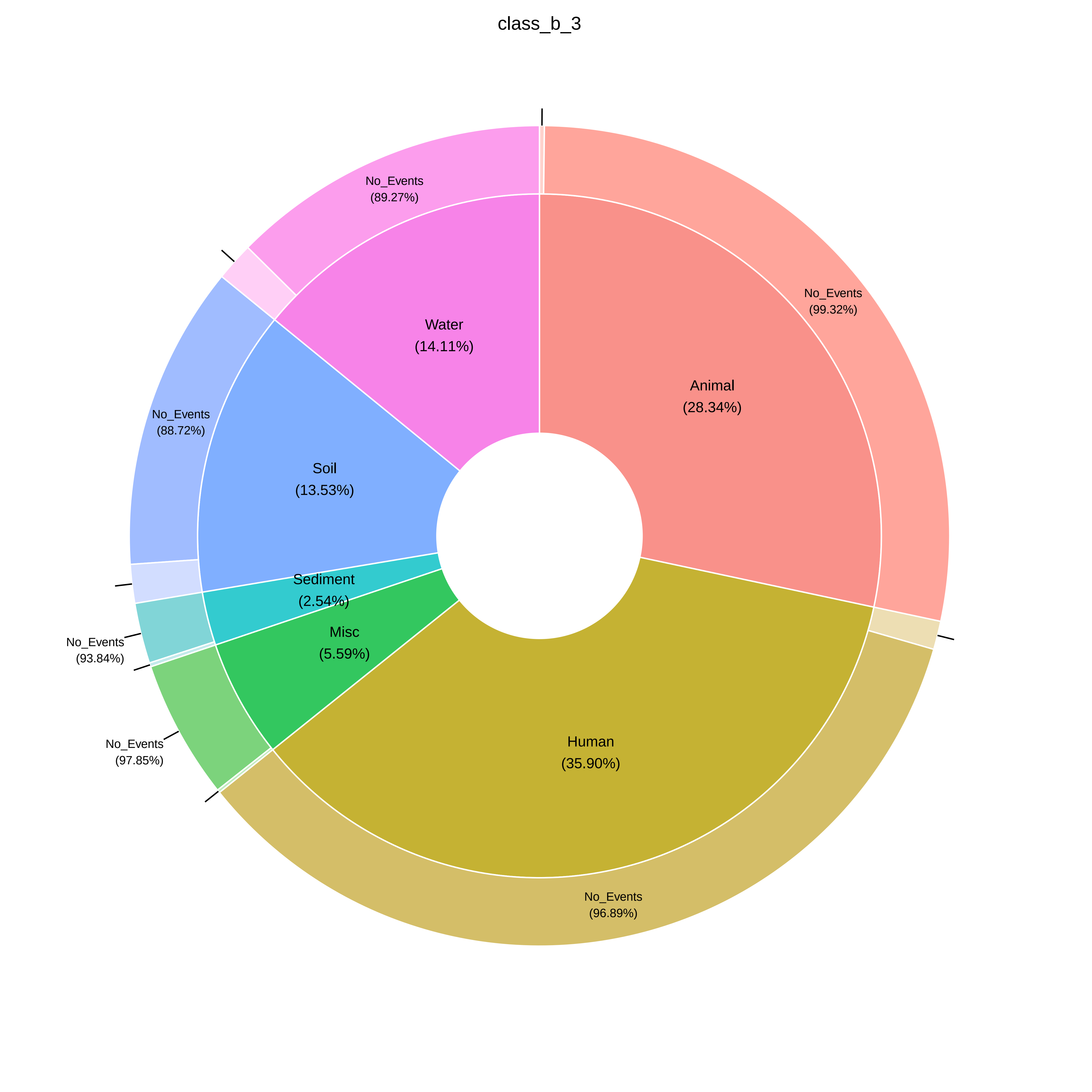
class_b_3
Animal
(28.34%)
No_Events
(99.32%)
Human
(35.90%)
No_Events
(96.89%)
Misc
(5.59%)
No_Events
(97.85%)
Sediment
(2.54%)
No_Events
(93.84%)
Soil
(13.53%)
No_Events
(88.72%)
Water
(14.11%)
No_Events
(89.27%)

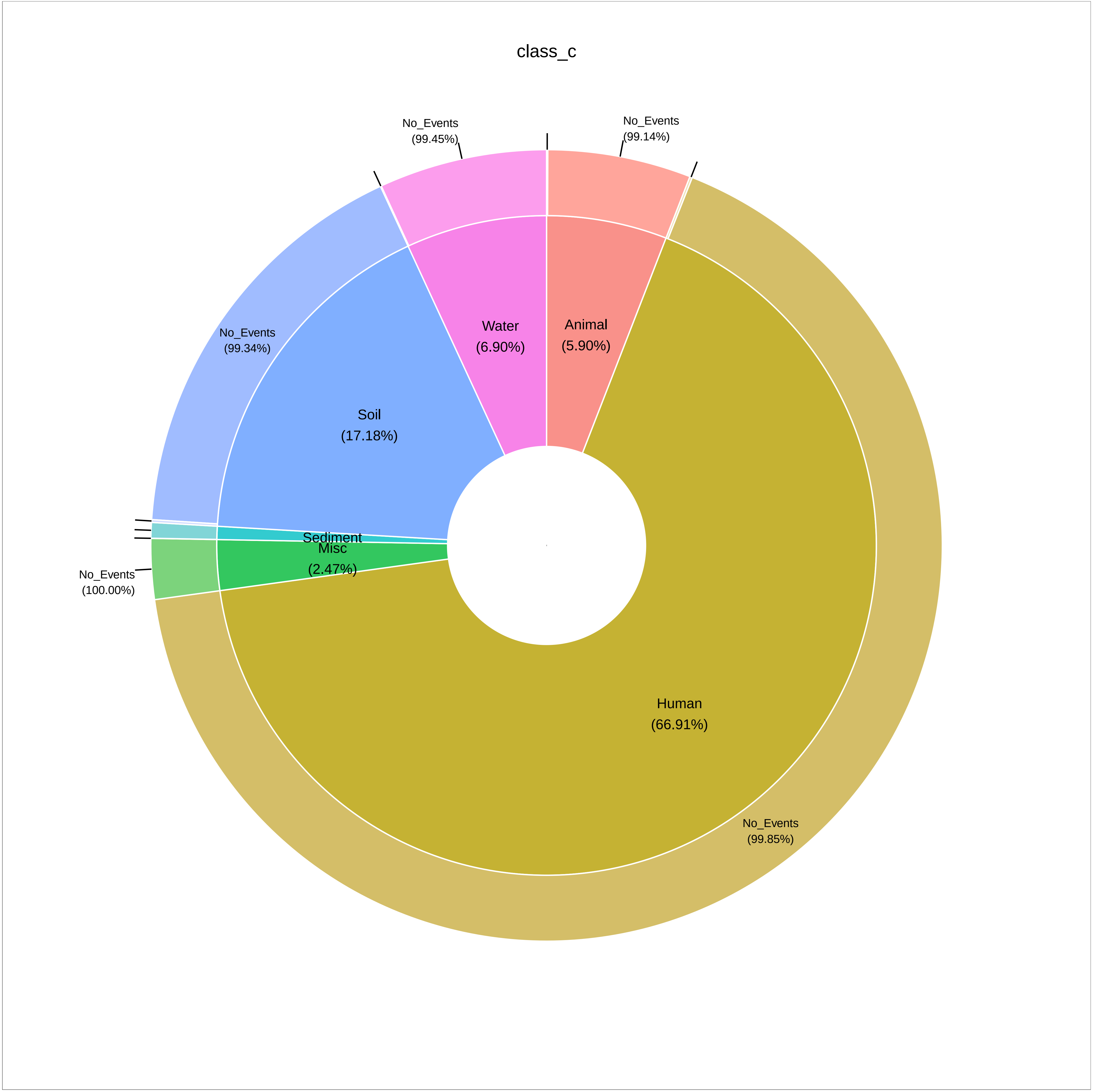

class_c
No_Events
(99.45%)
No_Events
(99.14%)
No_Events
(99.34%)
Water
(6.90%)
Animal
(5.90%)
Soil
(17.18%)
Sediment
Misc
(2.47%)
No_Events
(100.00%)
Human
(66.91%)
No_Events
(99.85%)

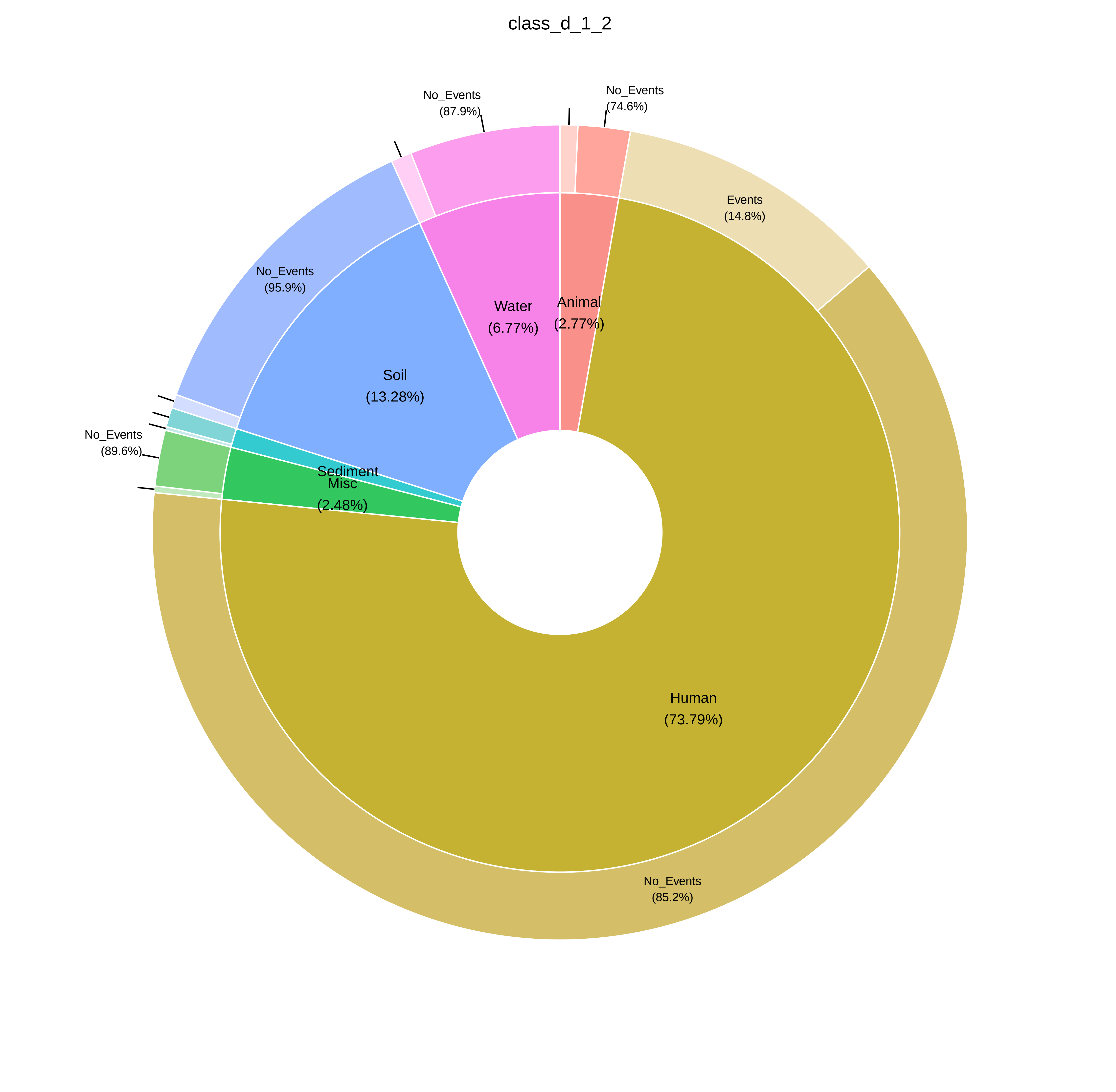

class_d_1_2
No_Events
(87.9%)
No_Events
(74.6%)
Events
(14.8%)
No_Events
(95.9%)
Water
(6.77%)
Animal
(2.77%)
Soil
(13.28%)
No_Events
(89.6%)
Sediment
Misc
(2.48%)
Human
(73.79%)
No_Events
(85.2%)

macrolide_phosphotransferases

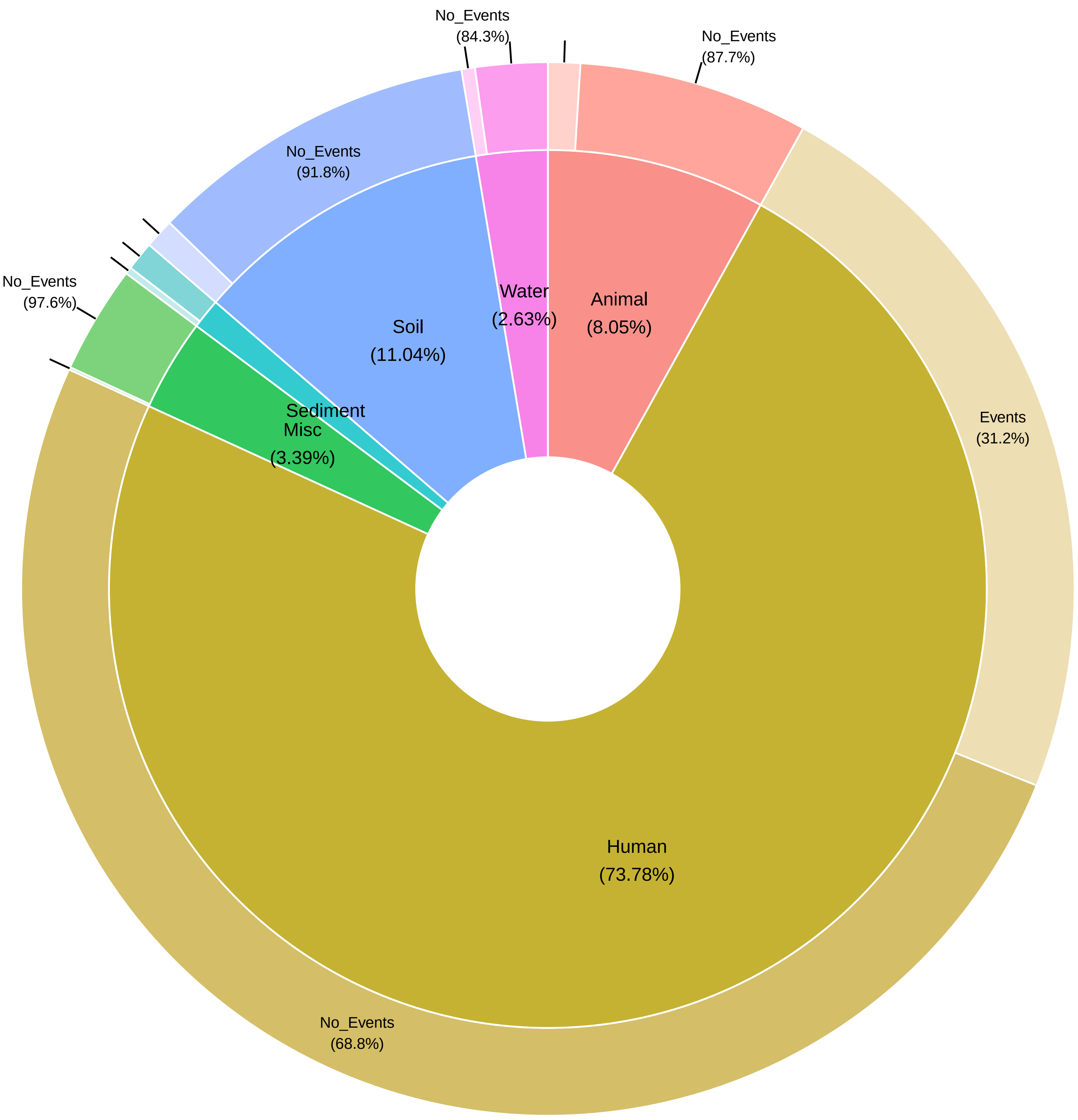

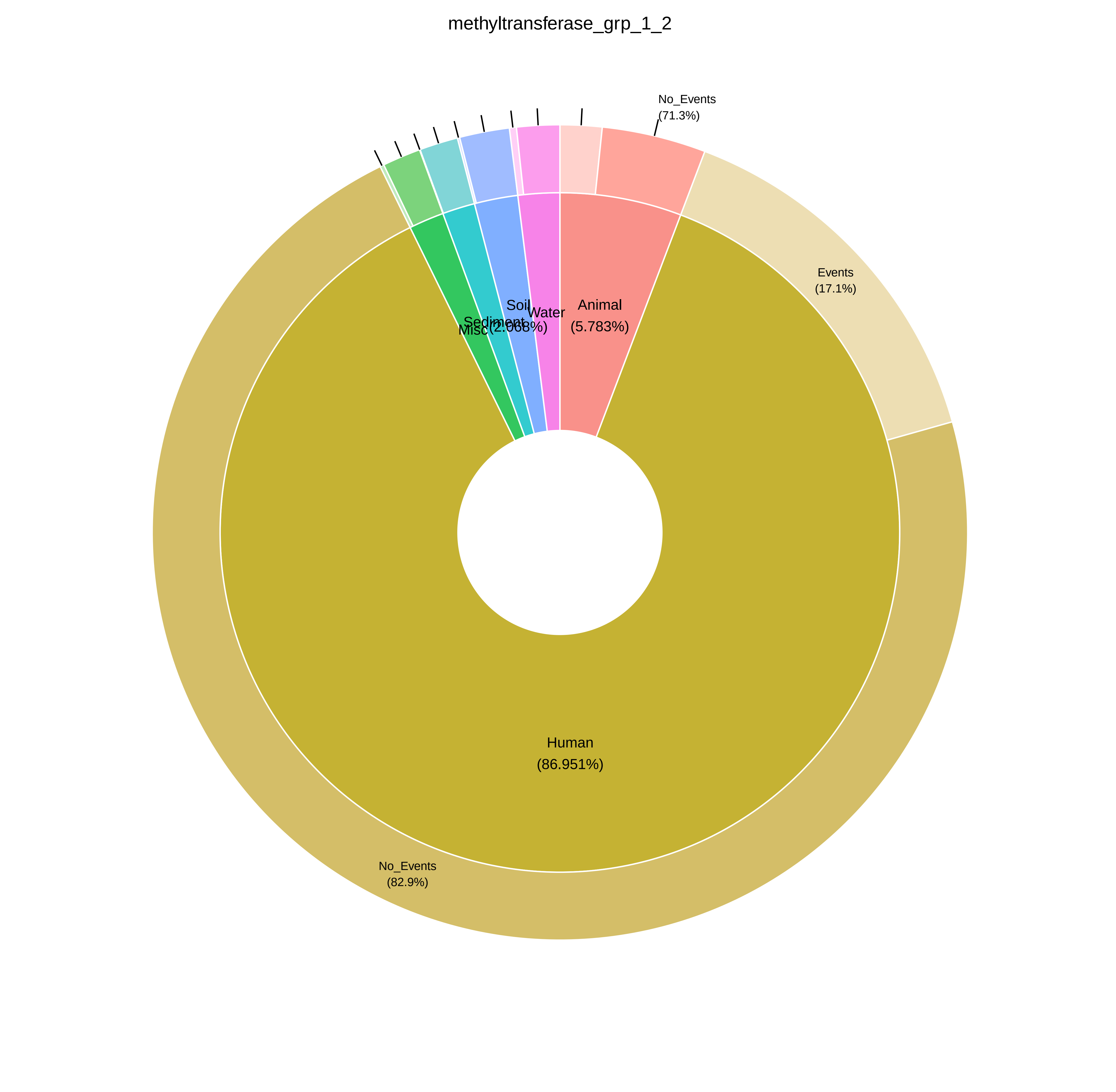

methyltransferase_grp_1_2
No_Events
(71.3%)
Events
(17.1%)
Animal
(5.783%)
Water
Soil
(2.068%)
Sediment
Misc
Human
(86.951%)
No_Events
(82.9%)

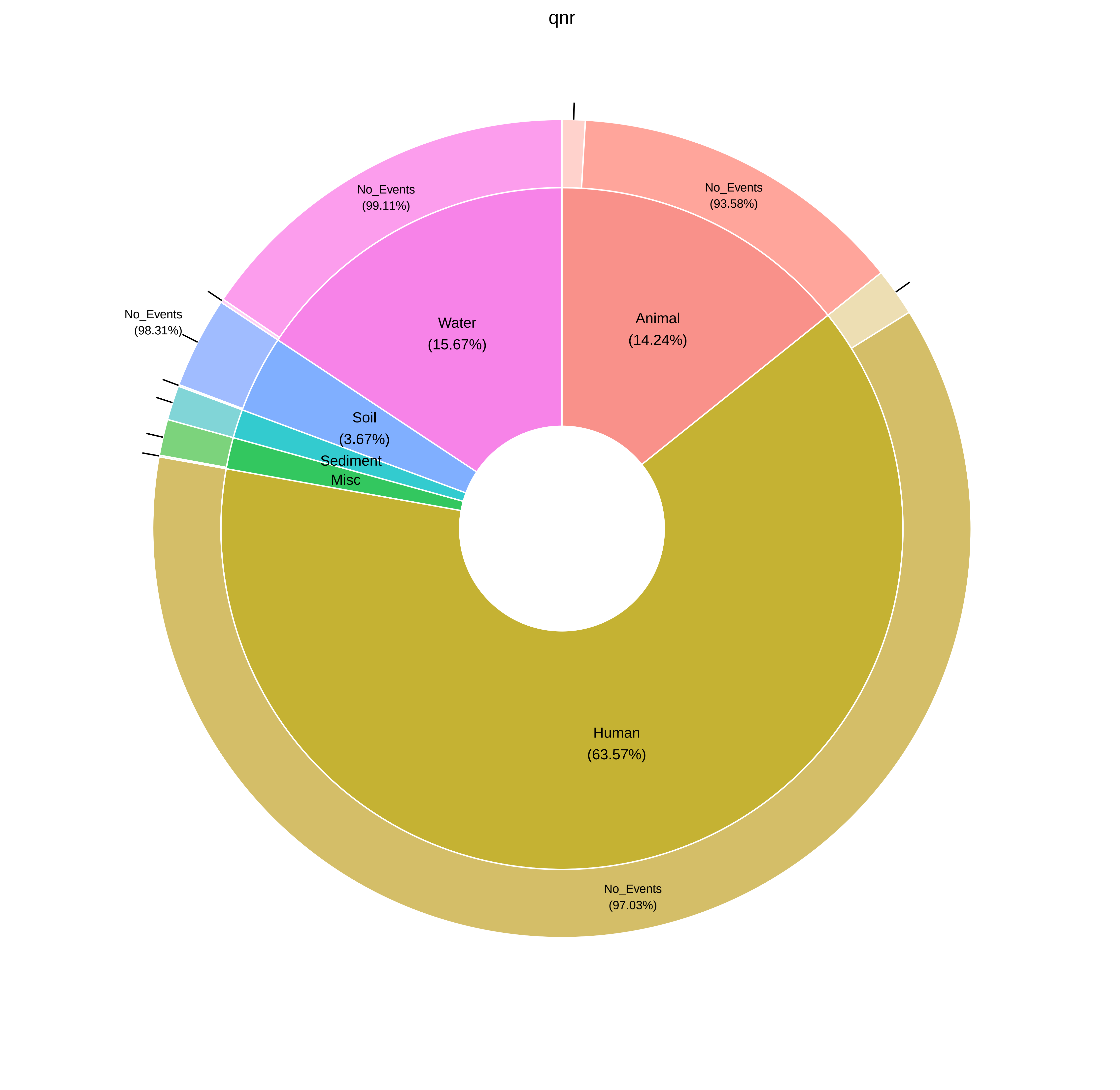
qnr
No_Events
(99.11%)
No_Events
(93.58%)
No_Events
(98.31%)
Water
(15.67%)
Animal
(14.24%)
Soil
(3.67%)
Sediment
Misc
Human
(63.57%)
No_Events
(97.03%)

tet_efflux

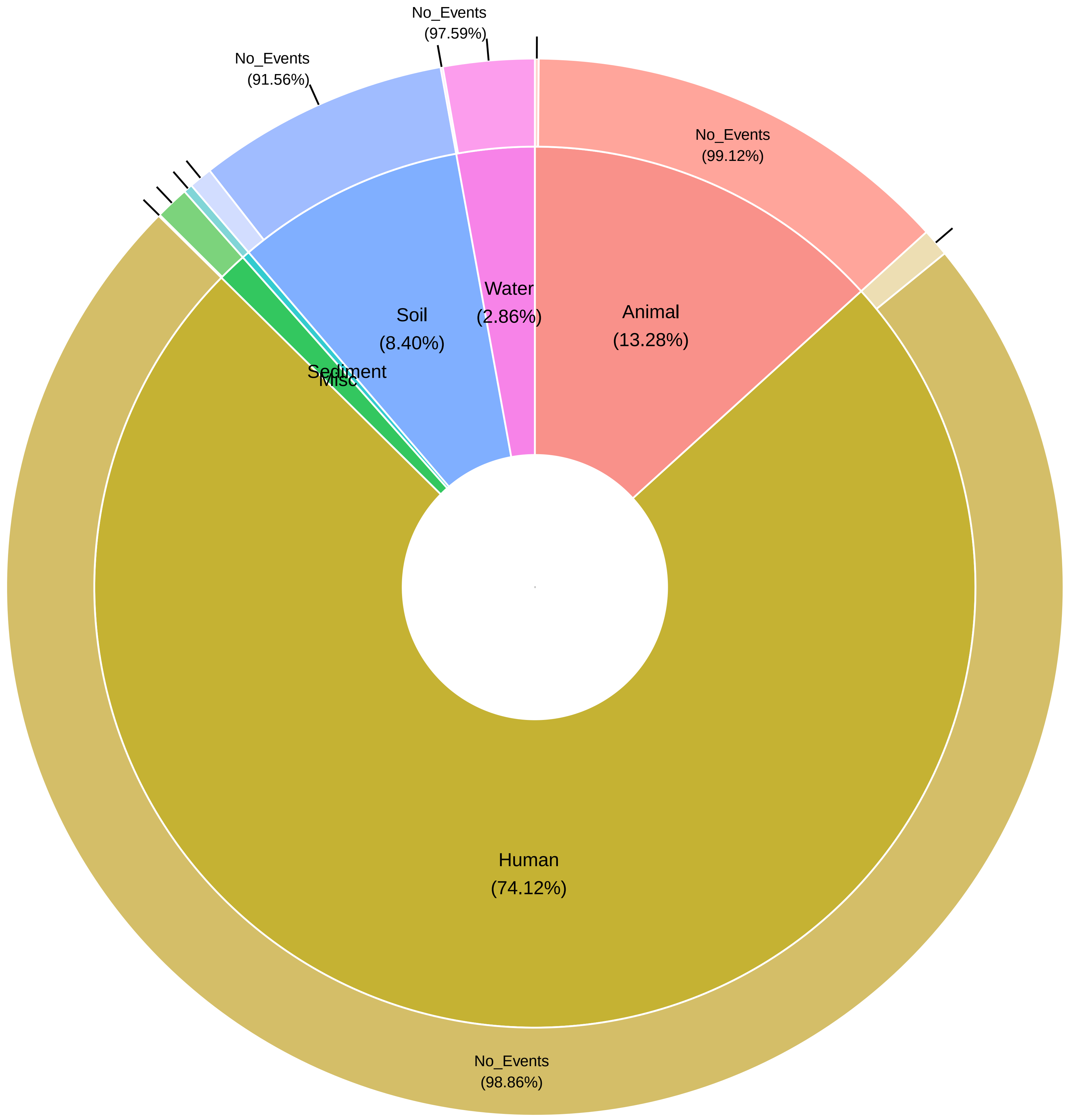

No_Events
(97.59%)
No_Events
(91.56%)
No_Events
(99.12%)
Water
(2.86%)
Animal
(13.28%)
Soil
(8.40%)
Sediment
Misc
Human
(74.12%)
No_Events
(98.86%)

tet_enzyme

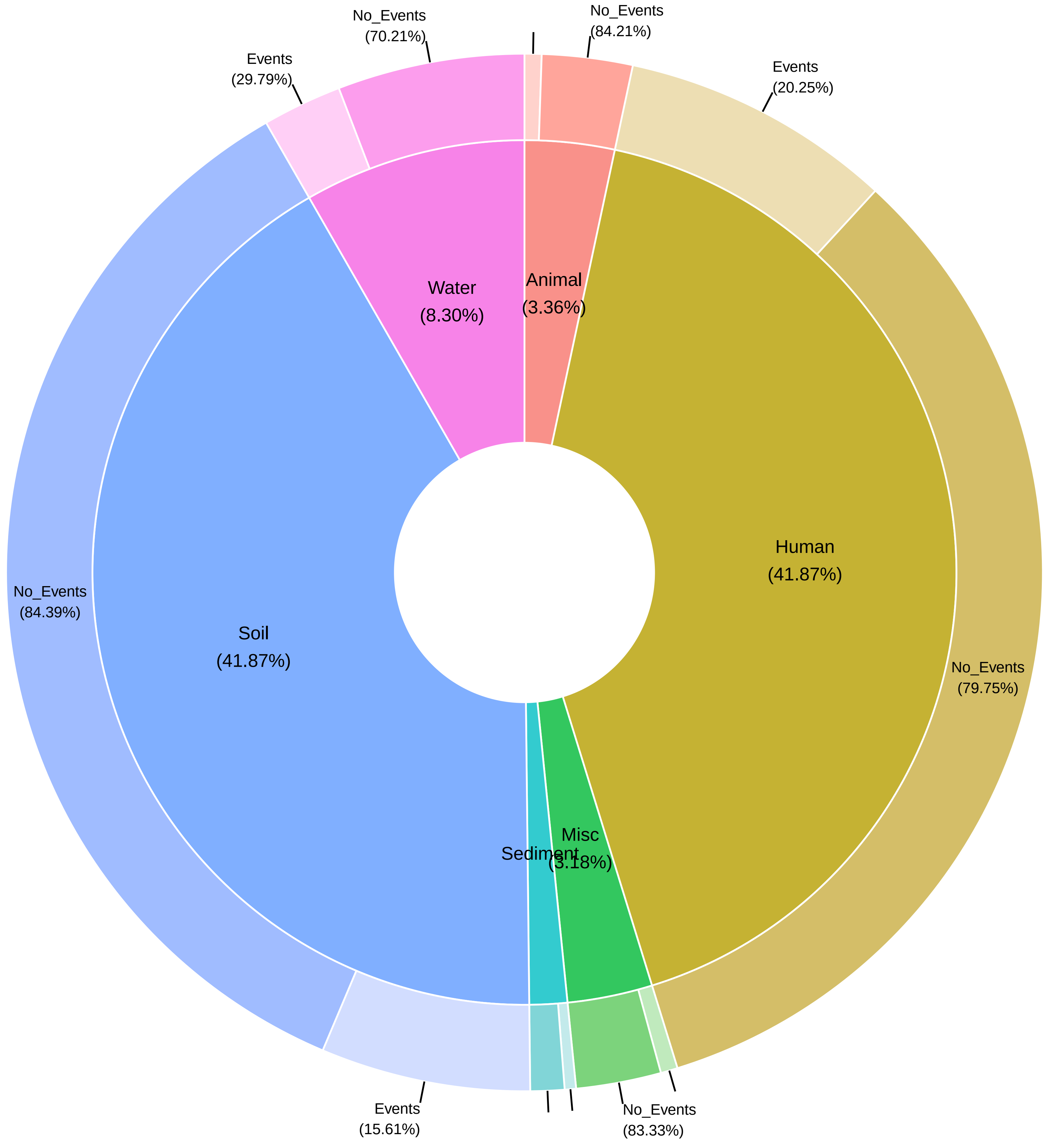

No_Events
(70.21%)
Events
(29.79%)
No_Events
(84.21%)
Events
(20.25%)
Water
(8.30%)
Animal
(3.36%)
Human
(41.87%)
No_Events
(84.39%)
Soil
(41.87%)
No_Events
(79.75%)
Misc
(3.18%)
Sediment
Events
(15.61%)
No_Events
(83.33%)

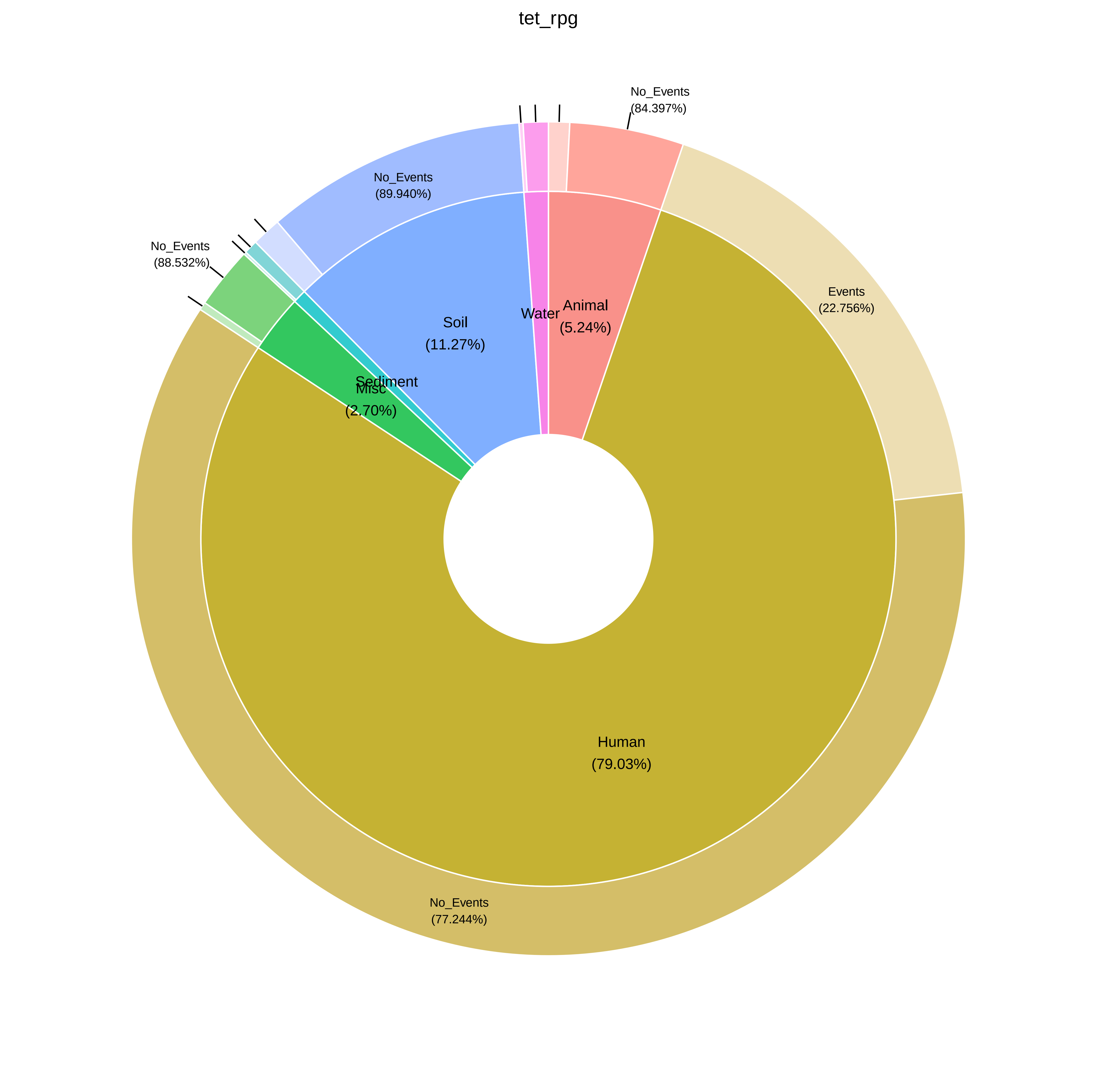
tet_rpg
No_Events
(84.397%)
No_Events
(89.940%)
No_Events
(88.532%)
Events
(22.756%)
Animal
(5.24%)
Water
Soil
(11.27%)
Sediment
Misc
(2.70%)
Human
(79.03%)
No_Events
(77.244%)
